# Evidence for climate-mediated range expansion of hybrid wood ants

**DOI:** 10.1101/2025.09.25.678544

**Authors:** Patrick Krapf, Patrick Heidbreder, Marjolein Bruijning, Patrick G. Meirmans, Sean Stankowski, Jonna Kulmuni

## Abstract

Climate change challenges many species. To persist, species can shift ranges, respond plastically, and adapt, which all require variation – often limited in natural populations. Hybridisation can quickly increase genetic variation, potentially facilitating climate adaptation, however it is unclear how hybrids respond to climatic changes. Here, we combine whole-genome, life-history, acute heat-shock, and climatic data of 69 wood ant populations across Finland to assess whether *F. aquilonia x F. polyctena* hybrids outperform their cold-adapted parent *F. aquilonia* in warming climates. Compared to *F. aquilonia*, hybrids are active and their offspring emerge earlier in spring, and they withstand acute temperatures relevant in nature better. Further, the hybrids’ border has shifted 200 km northwards, coinciding with the expansion of climatic conditions suitable for hybrids. Our results provide evidence that hybrids have an adaptive advantage over *F. aquilonia* due to their admixed ancestry, and recent climatic changes can lead to range expansion of hybrids.

## INTRODUCTION

Climate change is ubiquitous and challenges nearly every species. Since 1900, average global temperatures have drastically increased ^1^, with 2023 and 2024 being the hottest years since records began (Copernicus CS3). Concurrently, the frequency of extreme events such as heat waves, droughts, and heavy rainfall has dramatically increased in the last decades ^1,2^. The effects of higher temperatures and climate change are diverse ^3-5^ and are one of the major drivers causing global biodiversity losses ^6-8^. It is thus pressing and fundamental to understand and predict how species respond and persist under ongoing climate change.

Species can cope with changing climates through shifts in their habitat range, phenotypic plasticity, adaptation via genomic changes, or a combination thereof ^9^. Rapid range shifts are less likely for taxa with limited dispersal capacities, such as many plant and some animal species ^10,11^. Phenotypic plasticity allows individuals to adjust their phenotype to different environments within an organism’s lifetime and without an underlying genomic change ^12^. However, plasticity is finite, costly, and limited by the genomic makeup of an organism, and may constrain genomic adaptation ^13^ Hence, it is expected that many species will need to rely on rapid adaptation in a fast-changing world to persist ^14,15^.

Adaptation requires time and standing or novel genomic variation. While genomic variation may be present in large populations or populations with high mutation rates, many natural populations are small – often due to anthropogenic activities – and, as a result, have low levels of genomic variation ^16^. Hybridisation – defined as interbreeding between different populations or species ^17^ – has for long time seen as detrimental, but it can also provide a way to quickly increase genomic diversity ^18^, thereby increasing the adaptive potential of populations ^15^. For example, hybrid offspring of sunflowers *Helianthus annuus* and *H. petiolaris* can persist in extreme environments such as deserts or salt marshes while the parents cannot ^19^. Altogether, such elevated genomic and phenotypic variation in viable hybrids could make them better equipped to persist in rapidly changing environments ^20^.

To date, knowledge on the extent of hybridisation in natural populations is limited. Based on morphological data, the percentage of hybridising species was estimated to be approximately 10% in animals and 25% in plants ^21^. However, these estimates predate the advent of modern genomic analyses, and true rates are likely higher as hybrids frequently remain undetected ^21^. Irrespective of the exact hybridisation rates, they are expected to increase with natural and human-driven processes. For example, a recent southward move in an Atlantic puffin population led to the formation of a hybrid population in the Arctic ^22^. This is because a changing climate can cause range shifts ^14^ thus increasing the likelihood that previously isolated populations will interact ^23^. Besides increasing the chances of increased contact, a changing climate can also lead to the movement of taxa as seen in hybrid chickadees moving northwards ^24^. While these examples illustrate that hybrid populations are forming and moving in response to climate change, we lack understanding of whether hybridisation also promotes adaptation to climatic changes ^25^.

The *Formica* mound-building wood ants are a highly suitable group to study hybridisation and its implication for climate change adaption. This relatively young group has diverged into up to 13 species ^26^ within the last 500,000 years ^27^. Five of these occur in Finland with many of them hybridising naturally see below and ^28,29^, and hybrid populations account for a significant proportion of wood ant populations in Finland ^30^.

Wood ants are keystone forest species. They are frequently found in temperate and boreal forests where they provide important ecosystem services such as re-cycling nutrients ^31^, protecting the forest against bark beetles ^32^, and supporting over 200 other invertebrate species ^31^. The species differ in their social structures and dispersal abilities and also in their climatic adaptation. Three of the five species found in Finland are mainly single-queened and conduct long-range dispersal (*Formica lugubris, F. pratensis, F. rufa*), while the other two are multiple-queened and conduct short-range dispersal (*F. aquilonia, F. polyctena*) ^28,31^. However, the social structure and dispersal abilities do not reflect climatic adaptation: *Formica aquilonia* and *F. lugubris* are cold-adapted and are found at higher latitudes in Scandinavia and higher elevations in the European Alps. In contrast, *F. polyctena, F. pratensis*, and *F. rufa* are warm-adapted and found at lower latitudes in Scandinavia and medium to lower elevations in the European Alps ^33,34^. All wood ants also form characteristic and semi-stationary mounds, which can remain in the same location for multiple years ^31^. They thus need to respond to changing climates to ensure their persistence.

In Southern Finland, stable *F. aquilonia* x *F. polyctena* hybrid populations (as of now “hybrids”) are known ^20,29,30,35^. Recent simulation studies on *F. aquilonia, F. polyctena*, and these hybrid populations have shown that parental species have diverged from each other ∼224,000 years ago with gene flow mainly from cold-adapted *F. aquilonia* into warm-adapted *F. polyctena* ^36^. These simulations also suggest that hybridisation has been ongoing ever since. We assume that in Southern Finland, hybrids have been mating within hybrid populations. Also, two contemporary hybrid populations have formed in the early 1900s ^35^ and persisted ever since ^29^. However, we currently lack knowledge of whether other hybrid populations exist across Finland.

We define hybrids as individuals that display mixed ancestry between the parental species *F. aquilonia* and *F. polyctena* in the admixture analyses as well as their intermediate spatial placement between parental species in a PCA ^30^. These hybrid populations differ in some respects from their parental species. For example, hybrids are found in warmer and drier microclimates than populations of locally dominant *F. aquilonia* ^30^. Moreover, earlier results suggest that hybrids and *F. polyctena* perform significantly worse under acute cold shock than *F. aquilonia* ^37^. In contrast, hybrids withstand prolonged heat stress equally well as warm-adapted parent *F. polyctena* and significantly better than the cold-adapted parent *F. aquilonia* ^37^. However, data on acute heat shock are lacking. Also, hybrids harbour variation relevant for climate as selection favours alleles from the warm-adapted parental species in warm spring and alleles from cold-adapted parental species in cold springs ^37^. These findings suggest that hybrid wood ants may better cope with rapidly changing climate than parental species. However, extensive interdisciplinary analyses across space and time comparing hybrids and parents are needed to prove such broad-scale responses to a warming climate.

Here, we integrate whole-genome sequencing, life-history, acute heat stress, and climatic data on wood ant populations across Finland. This extensive and unique data set allows us to identify *F. aquilonia x F. polyctena* hybrid populations across Finland and compare hybrids with co-occurring cold-adapted *F. aquilonia* populations. We show that hybrids are frequently found in Finland, have a greater genomic diversity, which can lead to adaptive benefits. Hybrids are also likely to outperform *F. aquilonia* due to an earlier onset of mound activity and emergence of sexuals in the spring and an increased tolerance to acute heat stress (Fig. 1, top and middle panel). Consistent with these advantages, hybrid populations have persisted over 18 years, become slightly more abundant, and expanded their range to the north. The expansion of hybrid range coincides with expansion of climatic conditions suitable for hybrids over the past 18 years (Fig. 1, bottom panel). Overall, our data suggest that increasing temperatures due to climate change have facilitated a northward expansion of these hybrid wood ants. These findings highlight the major evolutionary and ecological role that hybrids can play in responding to a changing climate.

**Figure 1.**
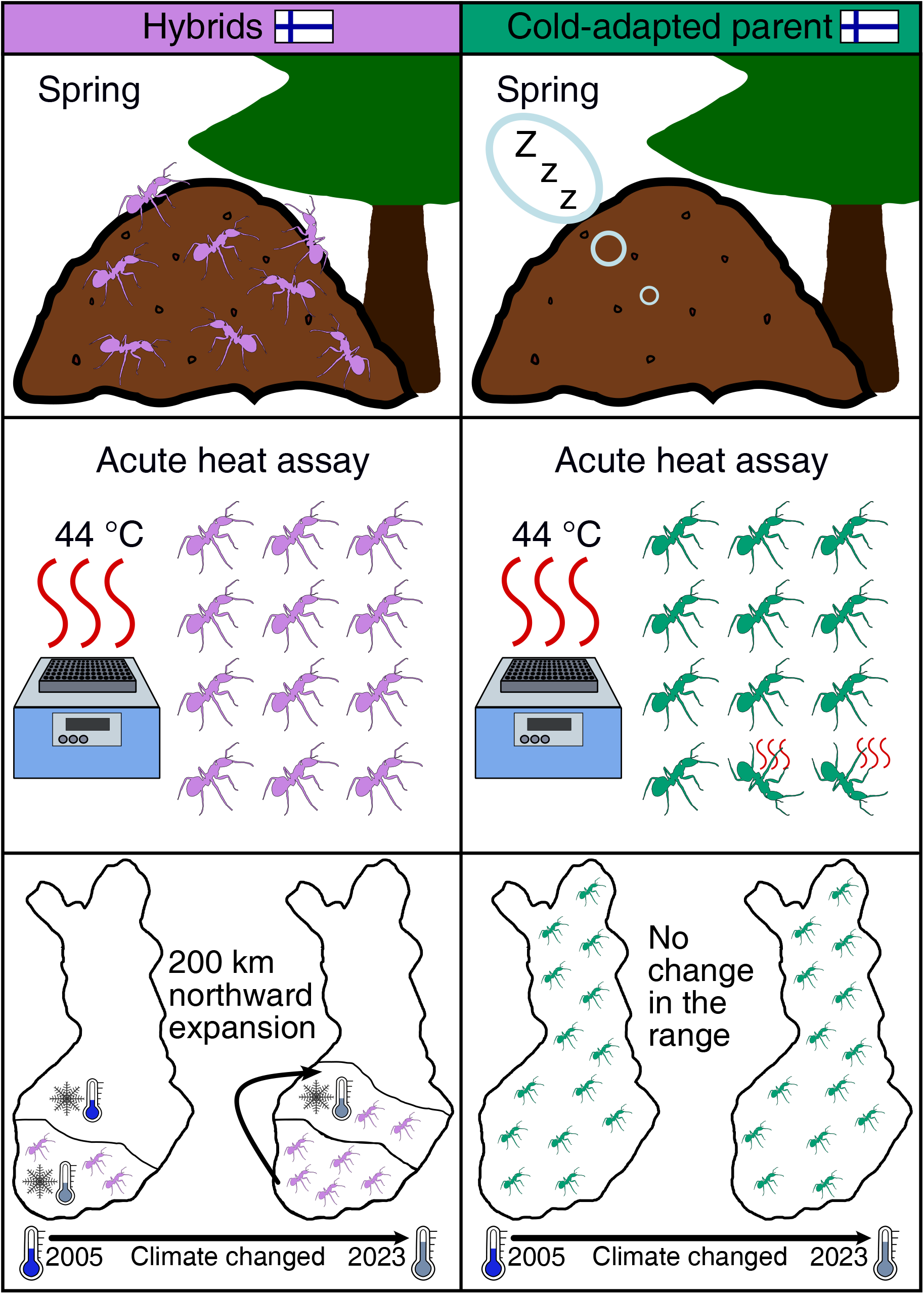
Compared to the cold-adapted parent *F. aquilonia, F. aquilonia* x *F. polyctena* hybrids are active earlier in the season, withstand acute heat stress better, and have expanded 200 km over 18 years in contemporary Finland. Top panel) Hybrids are already active in March and April, when *F. aquilonia* is still in diapause. Hybrids thus have fitness advantages when foraging food for queens at the start of the season. Middle panel) In acute heat assays, hybrids fall into heat coma at a higher temperature than *F. aquilonia*. Hybrids are thus likely able to continue foraging during hot periods in the summer, while *F. aquilonia* may forage less. Bottom panel) Over 18 years, hybrids have expanded 200 km northward coinciding with the expansion of climatic conditions suitable for hybrids over the past 18 years in Finland. In contrast, *F. aquilonia* did expand over time.

## RESULTS AND DISCUSSION

### F. aquilonia x F. polyctena *hybrids are among the most abundant wood ants in Finland*

Our first goal was to determine the distribution of *F. aquilonia, F. polyctena*, and their hybrids across contemporary Finland. To do this, we sampled 92 wood ant populations in 2023 (‘current sampling’) and used whole genome sequencing to identify species and hybrids. We found 48 *Formica aquilonia* (52% of all samples) and 21 *Formica aquilonia x F. polyctena* hybrid (23%) populations. In contrast to earlier morphological studies in Finland ^38,39^, we found no *F. polyctena* populations. We also detected other two *Formica* species and additional hybrids in the other samples with a relative frequency of <13% (Tab. S1-2, Fig. S1-2). We used whole-genome sequencing for species identification, and it remains a possibility that previous studies based on morphological identification have misidentified species.

We found hybrids more frequently than several wood ant species in Finland, and hybrids increased in frequency compared to previous studies. In this data set, 21 *F. aquilonia* x *F. polyctena* hybrid populations were found, which represents a 75% relative increase from the 12 populations observed in a previous study (‘previous sampling’) using a similar number of samples (N_2023_=21/92 (23%); N_2005- 2019_=12/87 (12%); Tab. S2; ^30^). This indicates that hybrids are persistent and one of the most abundant wood ant taxa in Finland. The ratio of hybrids compared to parental *F. aquilonia* or *F. polyctena* is high and similar or higher compared with other hybridising species pairs (some European ant species: 12-31%, ^40^; *Heliconius* butterflies: 7-13%, ^41^). For the remainder of this study, we focus on the comparison between *F. aquilonia* x *F. polyctena* hybrids and cold-adapted parent *F. aquilonia*.

Hybrids have a higher genomic diversity than their cold-adapted parent *F. aquilonia*. We calculated the observed heterozygosity using whole-genome data as a proxy for genomic diversity for samples found across the sympatric area (i.e. area where hybrids and *F. aquilonia* co-occur; Welch’s *t*-test: H_O_ hybrid: 0.11; *F. aquilonia*: 0.08; t =-4.70, df = 23.58, p-value<0.001; Fig. 2 A). These values are similar to the ‘previous sampling’ showing that hybrids have maintained diversity over time (H_O_ hybrid: 0.10; *F. aquilonia*: 0.08; ^30^). Such a higher genomic diversity can contribute to faster adaptation to climatic changes ^19,20^.

**Figure 2.**
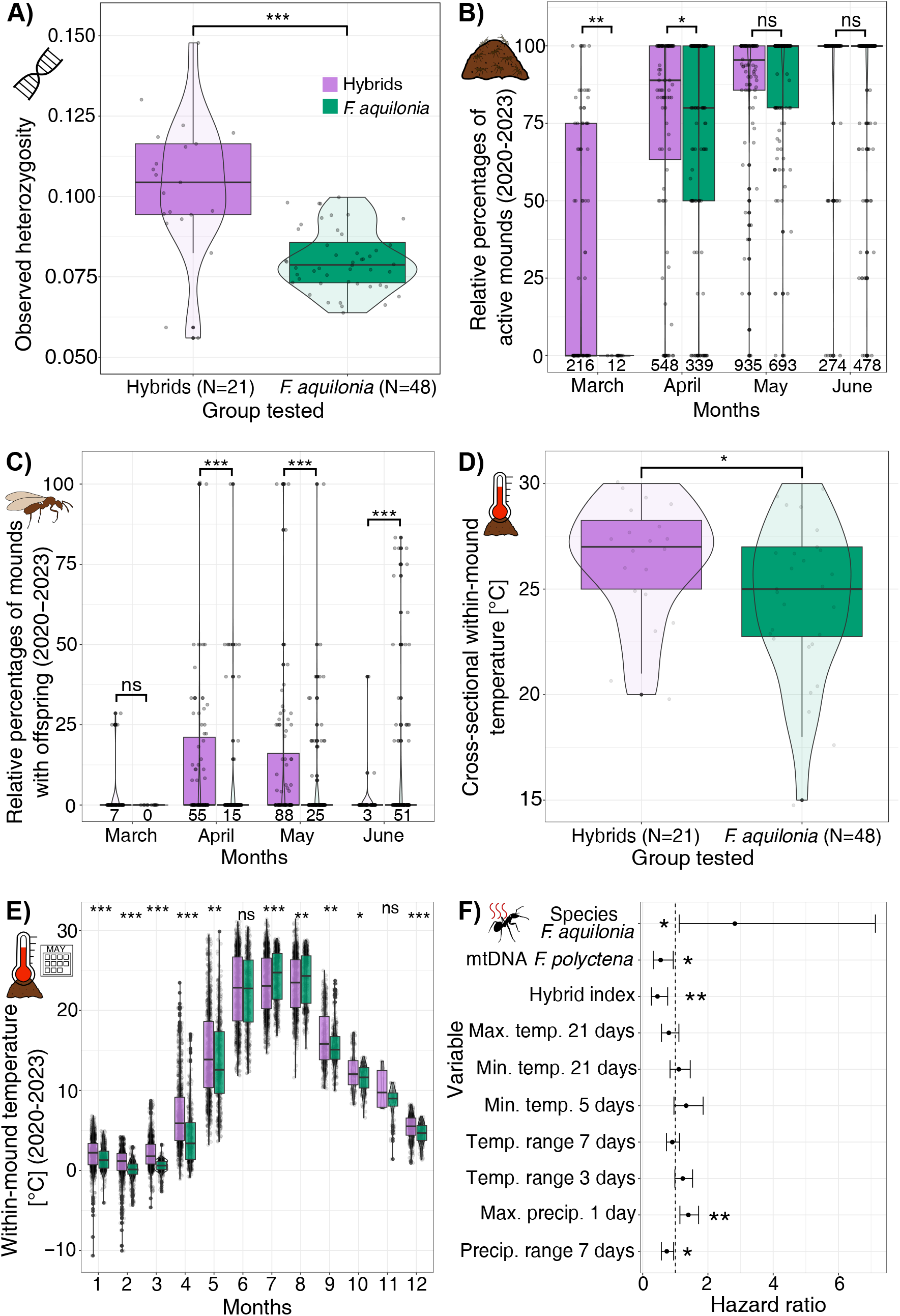
Recorded traits and characteristics for *F. aquilonia x F. polyctena* hybrids (purple) and *F. aquilonia* (green) populations. A) Observed heterozygosity including a mean comparison between hybrids and *F. aquilonia*. B) Relative percentages of active mounds in Southern Finland recorded between March and June for the years 2020 to 2023 for hybrids and *F. aquilonia*. C) Relative percentages of mounds with offspring in Southern Finland observed from 2020 to 2023. The numbers below the boxplots in panel B) represent the numbers of mounds checked for B) and C). The numbers below the boxplots in panel B) represent the numbers of mounds with offspring observed. D) Within-mound temperature measured at the top of the mound recorded during the sampling in the field in the year 2023. E) Within-mound temperature measured from HOBO loggers between 2020-2023 between hybrids and *F. aquilonia* (HOBO data points for hybrid mounds for each month over the three years = 837, 756, 837, 870, 961, 930, 961, 793, 383, 124, 8, 372; HOBO data points for *F. aquilonia* mounds each month over the three years = 341, 336, 372, 360, 480, 480, 496, 440, 293, 176, 76, 186). F) Expected hazard ratios to fall into heat coma presented for the species identity, mtDNA origin, hybrid index, and several weather variables prior to sampling. Significant comparisons are represented with asterisks, where * denotes p-value < 0.05, ** denotes p-value < 0.01, and *** denotes p-value < 0.001.

Further, we calculated the hybrid index and identified the mtDNA origin which showed that all *F. aquilonia x F. polyctena* populations are later-generation hybrids. The hybrid index ranged from 0.32 to 0.66 (i.e., scale 0 to 1, where 0 means only *F. polyctena* alleles and 1 means only *F. aquilonia* alleles; 359 ancestry informative markers; Fig. S3 A; ^35,42^) and indicates limited backcrossing. In nine out of 21 hybrid populations (43%), mtDNA originated from cold-adapted *F. aquilonia* mothers, and in 12 (57%) from warm-adapted *F. polyctena* mothers, their frequencies thus being indistinguishable from 50:50. This result differs starkly from the ‘previous sampling’, which identified ten *F. polyctena* maternal lines (83%) but only two *F. aquilonia* maternal lines in hybrids (17%; ^30^). For the hybrid samples, we also calculated pairwise mtDNA differences between the maternal lineages originating from both parental species. The mutations ranged from 76 to 86 mutations (Fig. S3 B) which corresponds to previous results ^30^.

### *Hybrids differ in several characteristics from the cold-adapted parent* F. aquilonia

Collected long-term data on the activity of ants and the emergence of sexual offspring show that hybrids are active earlier in the season, and their offspring emerges earlier and in greater numbers than *F. aquilonia* offspring. This data was recorded in three hybrid and three *F. aquilonia* populations from March until June from 2020 to 2023. We found that hybrid mounds were active earlier in spring than *F. aquilonia* mounds (Fig. 2 B, relative proportions calculated over all years; proportion test; March, X^2^ = 10.64, df = 1, p-value<0.001; April, X^2^ = 11.56, df = 1, p-value<0.001; Mai, X^2^ = 3.18, df = 1, p-value = 0.075; June, X^2^ = 1.71, df = 1, p-value = 0.191; corrected for multiple testing). Also, offspring from hybrid mounds emerged already in March and was significantly more frequent in April and May, compared with *F. aquilonia* mounds, where significantly more offspring emerged in June (Fig. 2 C; relative percentages calculated for observed mounds; two-sample test for equality of proportions with continuity correction; March, X^2^ <0.001, df = 1, p-value = 1.000; April, X^2^ =8.32, df = 1, p-value = 0.004; May, X^2^ = 19.87, df = 1, p-value<0.001; June X^2^ = 22.54, df = 1, p-value<0.001).

Further, cross-sectional and long-term data demonstrate that the within-mound temperature differs between hybrids and *F. aquilonia*. The within-mound temperature was warmer in the 21 hybrid mounds than in the 48 *F. aquilonia* mounds (measured from the mound top; hybrids mean = 26.38 °C; *F. aquilonia* mean = 24.45 °C, Fig. 2 D; Welch’s t-test, t =-°2.98, p-value = 0.019). Similarly, the long-term within-mound temperature measured from 2020 to 2023 (using HOBO data loggers) from six populations was significantly higher in hybrid mounds than in *F. aquilonia* mounds for almost the entire year, except for June and November (Fig. 2 E; Tab. S3). In contrast, within-mound temperature was significantly warmer in *F. aquilonia* mounds in July and August than in hybrid mounds.

The observed differences in spring activity are likely due to differences in diapause. Endogenous mechanisms are mainly controlling diapause in wood ants ^43,44^, but diapause is also affected by local adaption to climates ^43^. In contrast, temperature or photoperiod are expected to have little effect on diapause ^43,44^. Cold-adapted *Formica aquilonia* is likely genetically encoded to long winters ^34^ but also adapted to them thus having a longer diapause than, for example, warm-adapted wood ant species. Further, with climate change, the frequency of warmer winters is expected to increase, which may negatively impact *F. aquilonia* survival in future^45^: if winters are warm, Finnish *F. aquilonia* workers wake up during diapause thus using up fat reserves, which can lead to a reduced worker survival in subsequent spring ^45^. Hybrids, in contrast, with their genetic contribution of Southern European *F. polyctena*, seem to be encoded and adapted to shorter winters and having a shorter diapause. For example, in the Netherlands, *F. polyctena* mounds can already be active at the end of February (Krapf et al., unpubl.).

A shorter diapause in hybrids may represent a fitness over *F. aquilonia*. A shorter diapause and thus earlier activity allow hybrids to start foraging earlier in the season and to take advantage of longer spring and summer seasons. It further enables the earlier emergence of sexual offspring. In turn, hybrids may have a competitive and fitness advantage in obtaining nest sites and food as well over *F. aquilonia* and also other wood ants in Finland. Further, the within-mound temperature in wood ants is produced by workers that produce warmth with their metabolic heat ^46^ as well as due to their associated microorganisms ^47^. During the season, workers also continuously regulate the nest temperature by their metabolism, partial removal of mound material to reduce wall thickness, and movement of queens and brood inside the mound ^48^. Hybrids’ higher within-mound temperature, especially in the spring, could be an effect of differing worker behaviours, associated microorganisms or myrmecophiles, and nest construction techniques. It could further be coupled with the endogenous control of the diapause and contribute to an earlier activity in spring contributing to a competitive advantage of hybrids over *F. aquilonia*.

### Hybrids perform better under acute heat stress

Given that hybrids’ mounds are warmer, and mound surface temperature can reach up to 66 °C and higher (Photograph S1, Krapf et al., unpubl.), we compared the performance of individual hybrid and *F. aquilonia* ants in experimental heat stress assays. We tested 205 ants across 18 populations in a Critical Thermal Maximum assay (CT_max_; ^49^). In brief, we transferred ants to a 1.5 ml Eppendorf tube sealed with cotton and stored the tubes in a block heater (Eppendorf ThermoMix, Hamburg, Germany). We then increased the temperature from 40 to 49 °C in 20 minutes (Fig. S4A inset) and recorded ants’ status (active or knocked down) every minute, thus simulating semi-natural hot conditions.

Species identity, mtDNA origin, hybrid index, and weather characteristics prior to sampling affect the survival of ants in a CT_max_ assay. We conducted at Cox proportional hazard model to integrate heat data with species identity, parental mtDNA status, hybrid index, and weather characteristics prior to sampling (e.g., minima or maxima for temperature precipitation; overall model significance: LRT = 29.39, df=10.14; p-value = 0.001). *Formica aquilonia* has higher 2.8-fold higher estimated hazard to fall into heat coma earlier compared with hybrids (Fig. 2F, Tab. S4). In contrast, hybrids ants with *F. polyctena* mtDNA and higher admixed ancestry proportions had an 45% and 55% lower estimated hazard, respectively, to fall into heat than ants with an *F. aquilonia* mtDNA. The precipitation one day prior to sampling increased the hazard of falling into heat coma earlier by 40% in hybrids and *F. aquilonia*, while an increase in the range of precipitation seven days prior to sampling reduces the hazard of falling into heat coma earlier by 26%. Further, hybrids withstand acute heat significantly better than *F. aquilonia* (44.0 °C; X^2^ = 7.18, df = 1 p-value = 0.004; asterisk in Fig. S4 B, Tab. S5). This temperature is also close to wood ants’ thermal maximum at ∼45 °C.

Within hybrids, we found no difference between warm-or cold-adapted maternal mtDNA origin or the hybrid index (as a proxy for the contribution of warm-adapted alleles) in shaping acute heat tolerance. Hybrids with mtDNA of warm-adapted *F. polyctena* fell into heat coma at a similar temperature than hybrids with mtDNA of cold-adapted *F. aquilonia* (*F. polyctena*-mtDNA hybrids: 45.6 °C; *F. aquilonia*-mtDNA hybrids: 45.5 °C; one-sided Mann-Whitney-U-test, W = 856.0, p-value = 0.341; Fig. S4 C). Hybrids also withstand acute heat similarly well over the entire CT_max_ assay (Kaplan-Meier survival analysis, p-value = 0.40; Fig. S4 D). Also, hybrid index as a proxy for higher genomic contribution from warm-adapted *F. polyctena* did not influence hybrids’ performance under higher temperatures (linear regression, r^2^ = 0.01, F_1, 87_ = 1.901, p = 0.171, Fig. S4 E).

Our results demonstrate that hybrids perform better under acute heat stress than cold-adapted parent *F. aquilonia*. Our approach reflects naturally measured high temperatures in summer, where mound surface temperatures can reach over 66 °C. Previous results with a limited number of populations hinted that hybrids perform better under long-term heat exposure than *F. aquilonia* ^37^, but knowledge on acute heat exposure for populations across latitudes was lacking. Here, we significantly increased the sample size (from 3 populations each to N_hybrids_=21; N_*F*. *aquilonia*_=29) as well as geographic area of where the workers originated (from 86 to 460 km maximum linear distance between the samples).

We speculate that the hybrids’ mixed ancestry as well as the mtDNA of the warm-adapted parent *F. polyctena*, likely give hybrids an adaptive advantage over *F. aquilonia* in a warming climate. This is in line with recent results demonstrating that hybrid lineages can better cope with increasing temperatures than their parental species: for example, in the hybrid annual wildflower *Clarkia pulchella*, gene flow from a war-adapted species had a positive effect on fitness ^50^. Under warm conditions, F1 hybrid poplar trees survived better than parental species in a common-garden experiment ^51^.

One mechanistic hypothesis for the hybrids’ better performance under acute heat stress is that hybrids with warm-adapted ancestry may more effectively express heat-shock proteins (HSPs) than cold-adapted *F. aquilonia*. This has been observed in other warm-adapted ant species ^52^. Specifically, HSP60 and HSP90 are candidate HSPs, which are important for heat resistance in foraging ants in *F. cinerea* ^53^. Alternatively, DNA methylations may also contribute to a higher heat tolerance in hybrids as observed in the damselfly *Ischnura elegans* expanding to new ranges ^54^. Hybrids’ higher “baseline” temperature and heat stress resistance may enable hybrid workers to walk longer on hot surfaces, even if only for a few seconds. In turn, this may allow them to forage longer and more efficiently during hot summer days than *F. aquilonia*. However, this hypothesis needs to be tested in future studies.

Notably, also recent weather changes can affect survival under acute heat conditions. Our results demonstrated that changes in precipitation prior to sampling can increase or decrease ants’ performance.

This is in line with other studies showing that heat tolerance can be affected by desiccation and starvation in *Drosophila melanogaster* ^55^ as well as nutrition in ants ^56^. A hot and possibly drier climate can affect foraging behaviours in ants ^57^ and thus lead to different performance in heat tolerance assays.

### Hybrids have persisted and expanded 200 km northwards

To assess species and hybrid persistence over a total of 18 years, we compared the ‘current’ data sampled in 2023 to 44 populations from the ‘previous’ sampling. The ‘previous’ sampling included mainly samples collected between 2005-2008 with some additional populations sampled until 2019 that were already identified in Satokangas et al. (2023 ^30^). We found the same species or hybrids in 32 populations (73%) and species’ turnover in the other 12 populations (27%; Tab. S6). These revisited sites included ten hybrid populations, and we found that hybrids persisted in seven (70%) populations, some of these since 2005. Previous studies have indicated two isolated hybrid populations in Southern Finland have persisted for at least 19 years ^29^. One of these is estimated to have formed independently 13 to 47 generations ago, roughly at the beginning of 1900 ^35^. Here, we show, for the first time, that also other hybrid populations in Southern and Central Finland have persisted, at least 18 years.

The hybrid border has expanded northwards by ∼200 km in a timeframe of 18 years (Fig. 3 A). Data from the ‘current’ sampling (i.e., collected in 2023) demonstrated that, it was 1.6 times more likely to find hybrid populations at higher latitudes compared to the ‘previous’ sampling when using *F. aquilonia* as baseline (multinomial logistic regression; AICc = 238.30, Nagelkerke Pseudo-r^2^ = 0.38, odds ratio z-value =-4.39, p-value <0.001; delta AICc intercept-only model: 35.58; Fig. S4 F). In contrast, we found no evidence that the range of either parental species or other warm-adapted species has significantly changed (*F. aquilonia* no change; *F. pratensis* ∼145 m northwards shift, *F. rufa* ∼200 km southward shift; odds ratio z-value =-0.28, p-value = 0.078; Fig. 3 A). This indicates that the significant northwards shift is specific to hybrids. Further, this northward expansion is remarkable for such semi-stationary insects, as insects are expected to roughly move on average 12-15 km per decade ^11,58^.

**Figure 3.**
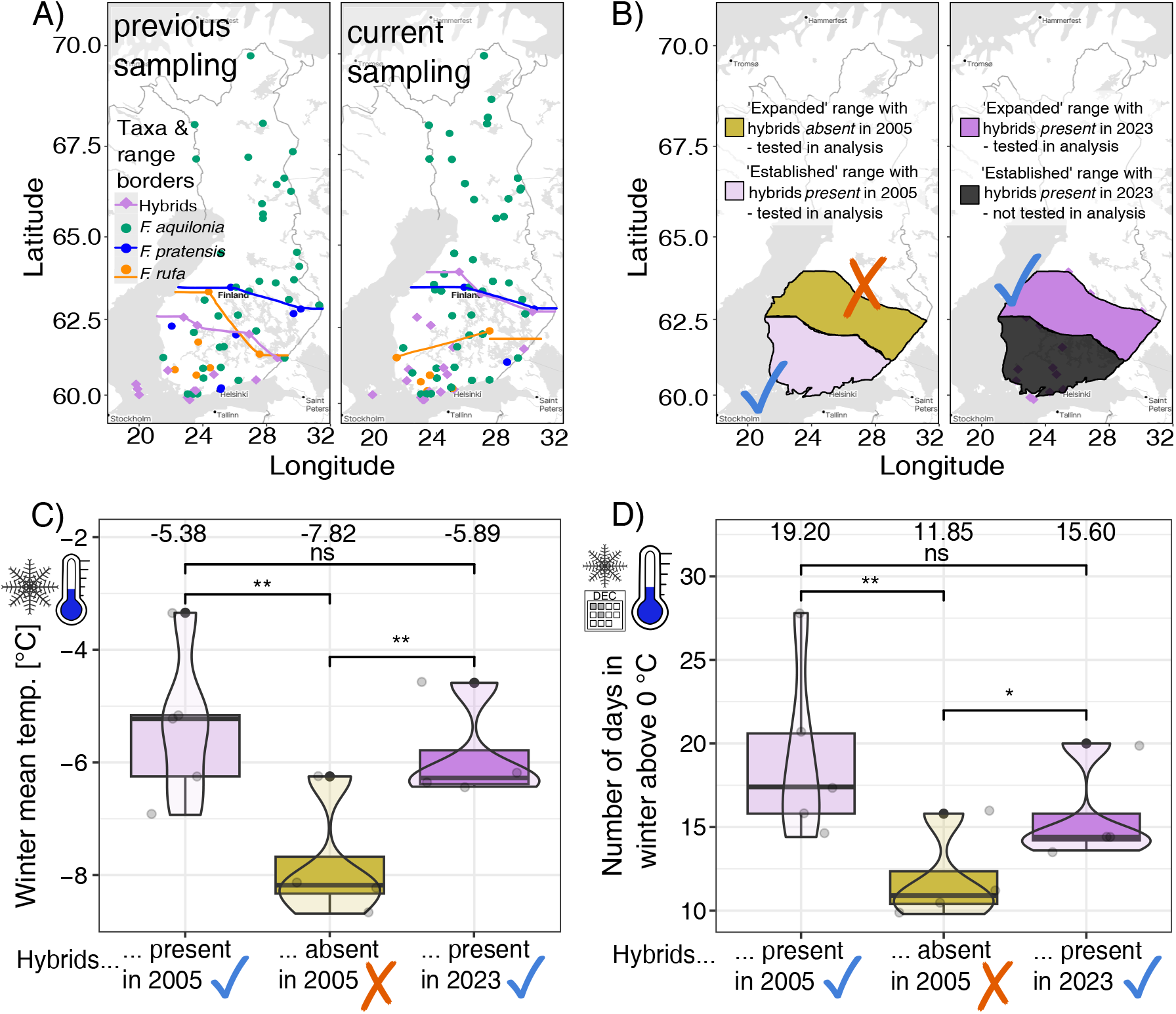
Sampling map and comparison of 5-year climatic averages of hybrid areas. A) Current and previous sampling map including taxa assignment based on whole-genome sequence data to identify wood ant species and hybrids. The previous sampling map was modified from Satokangas et al., (2023) ^30^. The current sampling originates from this data set. The lines represent species. B) Map displaying the hybrid range and ‘expanded range’ where hybrids were found and which were used for the temporal comparison. In C) and D), 5-year average of climatic variables were calculated backward from the sampling year 2005 (period 1999-2004) and 2023 (period 2018-2022). For simplicity, we refer to them as 2005 and 2023. C) Winter mean temperature and D) Number of days in winter above 0 °C (warm winter days). ‘Winter’ is defined as the diapause period between December to February. Values above the boxplots represent mean values. Asterisks represent bootstrapped p-values between hybrid areas at higher and lower latitudes and over time (Tab. S7) with * <0.05, ** <0.01, and ***<0.001. N ‘hybrid’ range in 2005 = 5. N ‘expanded’ range in 2005 = 4. N ‘expanded’ range in 2023 = 4.

### The northward expansion of the hybrids coincides with expansion of climatic conditions suitable for hybrids

If climatic changes have allowed a northward expansion of hybrids, we expect that the climate has changed in a way that is favourable for hybrids. Specifically, we expect that the climatic conditions in the ‘expanded’ range in 2023 should be similar compared to the conditions in the ‘established’ range in 2005 (Fig. 3 B, light and dark purple areas). In contrast, the climatic conditions in the ‘expanded’ range in 2005 should be unfavourable and differ from the climate in the ‘established’ range in 2005 (Fig. 3 B; yellow vs purple areas). To test these predictions, we calculate 5-year averages of six climatic variables (i.e., winter mean temperature, spring mean temperature, winter mean precipitation, spring mean precipitation, winter number of days above 0 °C and spring number of days below 0 °C), thus also accounting for yearly environmental fluctuations. Note, we use the terms ‘2023’ and ‘2005’ for simplicity when referring to the five-year averages from 2018-2022 and 2000-2004). Here, we only used hybrid populations, assuming that hybrid populations could follow these environmental changes and that these local climates are suitable for hybrids.

Following our expectations, the winter in 2023 in the ‘expanded’ range at higher latitudes (with hybrids present) was similarly warm and had a similar number of days above 0 °C as the winter in 2005 in the ‘established’ range at lower latitudes (Fig. 3 C-D; Tab. S7). At the same time, and also in line with our expectations, winter in 2005 in the ‘expanded’ range (with hybrids absent) was colder and not suitable for hybrids. However, not all aspects of the climate in the ‘expanded’ range have changed in such a way that matches the hybrid niche. Specifically, in spring 2023, temperatures in the ‘expanded’ range were still lower, there were more frost days below 0 °C, and there was more precipitation than in the spring in 2005 in the ‘hybrid’ range (Fig. S5 A-D; Tab. S7).

We also assessed whether the climate has changed as a whole across both hybrid and *F. aquilonia* populations. For this, we compared the same six environmental variables described above within and between hybrid and *F. aquilonia* populations and between previous and current sampling. The climate has changed similarly in both hybrid and *F. aquilonia* populations and followed the predicted climatic changes, with warmer and wetter winters and springs. Notably, these climatic changes were more pronounced in hybrid populations than in *F. aquilonia* populations (Fig. S6 A-E, Tab. S8). These results suggest both hybrids and *F. aquilonia* will need to cope with rapid changes as climate warming continues.

From these results, we can conclude that the climate appears to have become more favourable for hybrids at higher latitudes in Finland, but not in all respects. The increased winter temperatures follow our expectation and indicate that warmer winters may be more important for hybrids to expand northwards and persist than cold spells in spring, and that cold spells do not limit their persistence at higher latitudes. This rationale is supported by a study on *Aphaenogaster* ants demonstrating that increasing minimum temperatures along an elevation gradient in North America allowed the ant *A. rudis* to move upward in elevation ^59^. In general, warming temperatures under climate change can allow organisms to expand to new ranges ^11^. However, besides temperature also other factors such as precipitation or foraging strategy can also promote range shifts ^60^. For Finland, the Baltic Sea may have also contributed to the persistence and expansion of hybrids. It allows for mild winters on the Finnish coastal area, which had, for example, also a positive effect on the number of bird species in Southern Finland in the last decade ^61^.

While we have tested acute heat shock and calculated warm temperatures in winter and spring, also minimum temperatures can affect the expansion of ants ^62^. We have focused on CT_max_ rather than CT_min_ because hybrids were less cold resistant than parental species *F. aquilonia* ^37^. Further, because climate is warming ^1^ and CT_max_ is less plastic than CT_min_ ^63^, we were focusing on CT_max_ but acknowledge that testing CT_min_ may be interesting to test in future studies.

### Smaller mound volume and less divergent mtDNA at higher latitudes may indicate a recent northward expansion of hybrid populations

If hybrid populations have initially formed in Southern Finland and have recently expanded northwards or if the northward expansion is due to new hybridisation events, we expect hybrid mounds at higher latitudes to be smaller (as ants increase mounds volume over time). We also expect hybrid mtDNA in the expanded range to be less diverse than at lower latitudes since recently expanded hybrids had less time to accumulate mutations.

We found that hybrid mounds are significantly smaller at higher latitudes than *F. aquilonia* mounds, which are bigger but not significantly (hybrids: 9085.7 South and 5235.4 North; Welch’s t-test, t = 1.96, df = 17.33, p-value = 0.033; linear regression: r^2^ = 0.02, estimate =-1009.40, F-statistic = 1.32, df = 1 and 18, t-value =-1.15, p-value = 0.265; Fig. S7 A; *F. aquilonia* means: 8191.9 South and 12112.3 North; Mann-Whitney-U-test, W = 196, p-value = 0.057; linear regression: r^2^ = 0.08, estimate = 875.10, F-statistic = 4.77, df = 1 and 45, t-value = 2.19, p-value = 0.034). Notably, hybrid and *F. aquilonia* mounds do not differ significantly in their size regardless of whether they are found in the sympatric range or across entire Finland (Fig. S7 C-D).

Further, we also found that mtDNA of hybrids is less diverse at higher latitudes than at lower latitudes (i.e., fewer pairwise mutations in the mtDNA within the taxa; linear regression, hybrids: r^2^ = 0.03, estimate =-0.9825, F-statistic = 17.91, df = 1 and 570, t-value =-4.23, p-value<0.001, Fig. S7 B). In *F. aquilonia*, we found the opposite trend with a higher divergence at higher latitudes (linear regression; *F. aquilonia*: r^2^ = 0.03, estimate = 0.380, F-statistic = 211.70, df = 1 and 7308, t-value = 14.55, p-value <0.001).

The r^2^ values of both analyses are relatively small but may still indicate that hybrids have recently expanded northwards. The reduced hybrid mound sizes at higher latitude and the lower number of mtDNA mutations in hybrids may both indicate that they started building mounds at higher latitudes more recently. Further, the higher number of mtDNA mutations in *F. aquilonia* at higher latitudes may be a signal of lower population densities in Southern Finland marking the Southern range edge of *F. aquilonia*.

Previous work hinted that selection favours alleles from the cold-adapted parent in cold springs and alleles from the warm-adapted parent in warm springs ^*37*^. If such a signal is strong, hybrids may have a higher allele frequency of cold-adapted *F. aquilonia* alleles or *F. aquilonia* mtDNA origin at higher latitudes. However, both the hybrid index and mtDNA origin were not correlated with latitude (hybrid index linear regression, r^2^ =-0.02, estimate =-0.01, F-statistic = 0.36, df = 1 and 40, t-value =-0.50, p-value = 0.622, Fig. S7 E; mtDNA origin ANOVA, F-value = 0.979, df = 1 and 18, p-value = 0.336, Fig. S7 F). Nevertheless, we acknowledge that certain genomic regions and/or loci involved in the northward expansion and adaptation to cooler regions may not have been detected by simply looking at the ancestry proportion of the hybrid genomes.

While we present evidence for a climate-mediated range expansion of hybrid wood ants, it remains unclear how hybrids have expanded northwards. It could be that hybrids have initially formed in Southern Finland and then expanded northwards or if the northward expansion is due to new hybridisation events. Below, we speculate what may contributed to the hybrids’ expansion.

Hybrids seem to have similar polydomous colony structures as both parental species and thus probably also similar dispersal strategies. Both parents are poor at long-range dispersal via flights ^64^ and new mounds are usually established by budding (i.e., queens and workers found a new mound a few meters next to the original mound; ^65^). Budding thus allows a slow but continuous northward expansion but does not explain the recent 200 km expansion. However, males from both parental species disperse by flight ^66^, which could explain the northward expansion in hybrids – assuming that hybrids show the same behaviour. If this were true, we would expect to find backcrosses at higher latitudes, which we did not observe, making this scenario less plausible. Alternatively, hybrid populations may form via novel hybridisation events, but since *F. polyctena* is mainly absent from Finland (based on the genomic data here and in ^30^), this also seem unlikely. It may be that a sparse number of *F. polyctena* populations are present in Finland, which we have not found during our studies. This could allow new hybridisation events. Alternatively, it may also be that hybrids differ in their dispersal strategy from their parental species, and that both hybrids’ queens and males disperse by flight. Also, human-assisted dispersal of queens, which has been shown in a moving hybrid zone in *Solenopsis* ants ^67^ may lead to the establishment of hybrid populations at higher latitudes. These scenarios need to be addressed in future studies.

## Conclusions and broader implications

We combine whole-genome data of wood ants sampled in 2023 across Finland, with observational, experimental, cross-sectional and long-term environmental, as well as climatic data. Altogether, we found that *F. aquilonia x F. polyctena* hybrids have a higher genomic diversity than its cold-adapted parent *F. aquilonia*, hybrids are active with sexual offspring emerging earlier in the spring than in *F. aquilonia*. Hybrids also have an increased tolerance to acute heat stress due to their mixed ancestry and mtDNA origin of the warm-adapted parent. Hybrids are thus likely able to take advantage of longer spring and summer seasons and continued foraging capabilities during hot summer days suggesting a higher adaptive potential. Further, hybrids have persisted and expanded approximately 200 km northwards in the last 18 years; and this northward expansion of the hybrids coincides with extension of winter conditions suitable for hybrids. The persistence and northward expansion of hybrids at higher latitudes is probably the result of both climate change and the hybrids’ genomic composition and their corresponding shift in life-history traits. Future studies should investigate if the high hybrid prevalence, persistence, and expansion is also observed across other geographic regions and across the tree of life.

With ongoing climate change ^1^, hybridisation is expected to increase in frequency ^14^ due to an increase in secondary contact of species and/or populations. Out results suggest that hybrid wood ants with limited dispersal abilities may have competitive advantages over resident species in the face of changing climates similar as other organisms such as hybrid sunflowers ^19^. Hybrids are also able to track rapid changes in climate, which allows them to expand their distribution. They can thus be well-equipped to respond to or withstand climatic changes – possibly better than their parental species – as seen with these hybrid ants and in other hybrid organisms such as in oaks ^68^, birds ^24^, and fish ^69^. Further, wood ants as keystone species of boreal forests and support a wealth of biodiversity ^31^. Coupled with their abundance, wood ant hybrids play a major evolutionary and ecological role in the forest ecosystem. We thus argue that hybrids should also be considered in conservation planning ^70^, especially if they have a higher adaptive potential than parental species.

## RESOURCE AVAILABILITY

### Lead contact

- Requests for further information and resources should be directed to and will be fulfilled by the lead contact, Patrick Krapf.

### Materials availability

- This study did not generate new unique reagents.

### Data and code availability

- Raw whole-genome sequencing data have been deposited at NCBI project PRJNA1447136 and are publicly available as of the date of publication.
- Whole-genome and mtDNA VCF files and climate data have been deposited at Zenodo at 10.5281/zenodo.19563998, and are publicly available as of the date of publication. Additional code for whole-genome analysis is found under https://github.com/KraPat/WoodAnt_Species_and_Hybrid_Assignment
- Any additional information required to reanalyze the data reported in this paper is available in the supplementary material.

## ACKNOWLEDGMENTS

We thank Xavier Arnan for helpful feedback during the CT_max_ pilot studies, Benedikt Mader for help during field work, Peter Kuperus and Melanie Humphry for help during lab work, Anna Karrellos and Marit Kujit for help with sample handling, as well as Marit Kuijt, Noora Poikela, Beatriz Portinha, and Ina Satokangas for helpful discussions during data analysis and writing. We wish to acknowledge CSC – IT Center for Science, Finland, for computational resources. The maps were created using Stadia Maps (stadiamaps.com), Stamen design (stamen.com), and OpenStreetMap (openstreetmap.org/copyright).

This work was funded by the Suomen Akatemia (Research Council of Finland) grant no. 346805, 324485, and 353156 awarded to J.K., the University of Helsinki LUOVA Doctoral Programme funded Doctoral researcher position awardee to P.H., and the European Union’s Horizon Europe programme under Marie Skłodowska-Curie Actions (MSCA) - Postdoctoral fellowship grant agreement no. 101204375 awarded to P.K. The climate data used were downloaded from the Finnish Meteorological Institute, FMI (under https://paituli.csc.fi/home.html).

## AUTHOR CONTRIBUTIONS

Conceptualization: PK, JK

Methodology: PK, MB, PGM, SS, JK

Software: PK, PH, MB, PGM, SS, JK

Validation: PK

Formal analysis: PK,

Investigation: PK, PH

Resources: JK

Data curation: PK, PH

Writing - Original Draft: PK, JK,

Writing - Review & Editing: PK, PH, MB, PGM, SS, JK

Visualization: PK, JK

Supervision: JK

Project administration: PK, JK Funding acquisition: PK, JK

## DECLARATION OF INTERESTS

The authors declare that they have no competing interests

### DECLARATION OF GENERATIVE AI AND AI-ASSISTED TECHNOLOGIES

During the preparation of this work, the author used ChatGPT in order to help annotating R code. After using this tool or service, the author reviewed and edited the content as needed and take full responsibility for the content of the publication.

### SUPPLEMENTAL INFORMATION

**Document S1. Figures S1–S7, Tables S1–S8, and supplemental references. Photograph S1. Photograph of a mound and its surface temperature in plain sunlight during summer related to the section “*Hybrids perform better under acute heat stress*”**.

## STAR⍰METHODS

### EXPERIMENTAL MODEL AND STUDY PARTICIPANT DETAILS

#### Animals

This study used ants from a non-threatened and non-protected ant species in Finland. No license or permit was required to conduct these assays. Nevertheless, we followed the highest ethical standards and international law for ants collected. We also followed two of the 3Rs Principle: i) We reduced the number of workers collected to a minimum (Reduction principle) and iii) we created the best accommodation possible for ant workers (Refinement principle). Further, all wood ants used in the subsequent analyses were collected by the authors of this study. Species or hybrid identity is presented in Table S1.

## METHOD DETAILS

### Sampling and recording of mound and environmental data

From June to August 2023, *Formica* wood ant mounds were sampled from 92 populations across Finland (Fig. 3 A, map created using ggplot2 ^71^, ggpubr ^72^, and ggsignif ^73^ in R v4.3.0 and RStudio v2025.09.0+387). From each mound, approx. ten workers were collected alive for thermal tolerance assays (detailed below), and a handful of workers were collected and stored in 96% ethanol for genomic analyses. Whenever possible, workers from the same mound and species were collected in 2023 as used in Satokangas et al. (2023; ^30^) for genomic and ecological analyses. During sampling, we recorded the within-mound temperature and mound size for each mound. We measured within-mound temperature from the highest point of the mound using a core thermometer (Xavax 111381, Monheim, Germany; Tab. S1). We also measured the mound length, width, and height using a measuring tape and calculated the mound volume following ^74^ and using the equation: 2/3 × π × D/2 × d/2 × H where D is the length, d is the width, and H the height of the mound.

#### Genomic analyses

##### DNA extraction, whole-genome sequencing, and variant identification

To identify hybrids and different species, we extracted DNA from a single diploid worker per mound and population (N=98; after filtering N=92) using the Qiagen DNEasy Blood and Tissue kit (Qiagen, Hilden, Germany; N=81) following the manufacturer’s protocol or a modified CTAB extraction protocol (N=17; ^75^). In both procedures, DNA was eluted in 100 µl AE elution buffer from the Qiagen kit. Library preparation and whole-genome sequencing were outsourced to a commercial provider (Novogene, UK). Whole-genome sequencing was conducted on an Illumina NovaSeq X Plus with 150bp paired-end sequencing.

Raw Illumina reads and adapter sequences were trimmed using trimmomatic (v0.39; ^76^) before mapping against a Finnish *F. aquilonia* × *F. polyctena* hybrid reference genome ^77^ using bwa mem (v0.7.17; ^78^). Using picard tools (v2.27.5; http://broadinstitute.github.io/picard), duplicates were removed with the command “MarkDuplicates”, read groups were added with the command “AddOrReplaceReadGroups”, and bam files were indexed using “samtools index” in samtools v1.2.1 ^79^. The average read depth was 15.7×, as computed across individuals using mosdepth (v0.3.6; ^80^) after duplicate removal. Overlaps were clipped using “bamUtil” and the function “bam clipOverlap” (v1.0.15, https://github.com/statgen/bamUtil/releases).

##### SNP calling and filtering data

For single nucleotide polymorphism (SNPs) calling, we used these 98 samples (‘current’ sampling) plus additional 85 samples from a so-called ‘previous’ sampling from Satokangas et al. (2023) ^30^ as a reference. We called SNPs with the freebayes software v1.3.6 ^81^. To speed up the SNP calling, 50 kb regions of the reference genome were defined using the script “split_ref_by_bai_datasize.py” (available under https://github.com/freebayes/freebayes/blob/master/scripts/split_ref_by_bai_datasize.py; main settings: --k option to disable population priors; --genotype-qualities to write genotype qualities to a file; -- skip-coverage to skip high coverage regions; --limit-coverage 100 to only use regions with a maximum coverage of 100; --use-best-n-alleles 3 to only consider 3 alleles to reduce runtime). The resulting VCF file was normalised using bcftools norm-function (v1.10; ^82^). Using bcftools, sites located at less than two base pairs from InDels were excluded, as well as sites that were supported by only Forward or Reverse reads. Furthermore, we only kept biallelic SNPs with quality values equal to or higher than 30 and, we filtered sites based on individual sequencing depth: Genotypes were marked as missing when their depth exceeded twice the mean depth of the individual in question. Genotyping errors (e.g., misaligned reads) were removed by excluding sites that displayed heterozygote excess after pooling all samples using vcftools v0.1.17 ^83^. Individual genotypes with a quality of lower than or equal to 30 and a depth of coverage lower than or equal 8 were set as missing data. Sites with more than 10% of missing data over all samples were removed from the next steps. Lastly, singletons were removed (minor allele count <2) using vcftools.

After these filtering steps, 3,324,311 SNPs were kept from 179 individuals with sequencing depths of at least 8×, including 92 out of the 98 samples from the ‘current’ sampling in 2023. Using vcftools, the final dataset was thinned with 20 kb distances to minimise linkage disequilibrium. We further excluded chromosome three (the social chromosome with reduced recombination due to supergene inversions; ^84^) as well as chromosome ‘zero’, which contains all unscaffolded contigs with no known genomic location ^77^. The final data set resulted in 9,979 genome-wide SNPs used for the genomic analyses described below.

#### Species and hybrid identification

##### Principal component analysis (PCA)and admixture analyses

To assess the species’ identity, genomic structure and hybrid status, a principal component analysis (PCA) was conducted in plink (v2.00a6.12, ^85^) using the nuclear genomic SNPs data from all samples. We expected that genomically diverged species and hybrids would form separate clusters in the PCA.

To assess the extent of admixture and thus hybridisation in the samples, we performed an admixture analysis ^86^ with the 9,979 nuclear SNP data. For admixture, the number of clusters K was chosen to range from K = 2 to 10, without prior knowledge of the taxa but including samples and species identity from the ‘previous’ sampling ^30^. The most likely value of K was chosen based on the lowest cross-validation error rates. We further extracted the allele frequencies from the most likely K to be used in statistical analyses.

##### Mitochondrial haplotype network

To create a haplotype network using the mitochondrial DNA only, we called SNPs from the mitochondrial genome using all 179 samples. We used the same filtering steps as with the nuclear DNA. After filtering, 466 SNPs were kept. A consensus sequence for each sample across all samples was created by converting the VCF file into a fasta-file using the faidx function in samtools. Next, the fasta-file was converted to a phylip-format and used to create a median-joining network in PopArt ^87^. Species that share identical or similar mitochondrial haplotypes should cluster more closely together in the network.

##### Selecting hybrids and F. aquilonia for subsequent comparisons

After reliably identifying species and hybrids, we only used samples from *F. aquilonia x F. polyctena* hybrids and its cold-adapted *F. aquilonia* parent for subsequent analyses. We did not find the warm-adapted parent *F. polyctena* in the data set. We also excluded the warm-adapted *F. rufa* and *F. pratensis*, as well as cold-adapted *F. lugubris* because they are either single-queened (or facultatively have only a few queens) and not as dominant locally as *F. aquilonia* or hybrid. Both *F. aquilonia* and hybrids are also socially more similar (i.e., both are multiple-queened and form supercolonies; ^88^).

#### Downstream genomic analyses

##### Observed heterozygosity, hybrid index and intraclass heterzoygosity calculation

To assess whether hybrids had a greater genomic diversity than *F. aquilonia*, we first calculated the observed heterozygosity for each sample from the current sampling using vcftools and then tested if their means differed significantly using a Welch’s t-test. To calculate the hybrid index for the hybrid samples, we used the ‘previous’ and ‘current’ sampling and filtered the VCF-file to only keep the parental species *F. aquilonia* and *F. polyctena* and the *F. aquilonia* x *F. polyctena* hybrids. Then, we removed monomorphic sites, selected ancestry informative markers (AIMs) with an allele frequency difference of 0.95 between *F. aquilonia* and *F. polyctena* ^77^, and filtered the data to keep only markers with less than 5% missing data. Lastly, we thinned the VCF file to 1 SNP/500 kb yielding 359 SNPs. We then exported the VCF file and calculated the hybrid index for each individual as following: sum of *F. aquilonia* alleles / ((sum of homozygous *F. aquilonia* and homozygous *F. polyctena* alleles) - 2 * number of missing genotypes). Using the hybrid index and individuals from the ‘current’ sampling 2023, we conducted a linear regression testing whether the hybrid index is associated with the latitude as well as a linear regression testing whether it is associated with the temperature when ants fall into heat coma (section “*Critical Thermal Maximum (CT*_*max*_*) assays*”). We also calculated the intraclass heterozygosity for each individual as following: sum of heterozygous alleles / (sum of homozygous *F. aquilonia* alleles, sum of heterozygous alleles, and sum of homozygous *F. polyctena* alleles).

##### Identifying mtDNA origin and pairwise mtDNA mutations

Using the mtDNA haplotype network, we identified the maternal species from which hybrids originated (i.e., *F. aquilonia* or *F. polyctena*) as mtDNA is maternally inherited and calculated pairwise mtDNA mutations for each sample pairs. Using an Analysis of Variance (ANOVA), we tested whether the mtDNA origin is associated with the latitude. Further, we correlated the pairwise mtDNA mutations for the hybrid maternal lineages against the geographic distance using a linear regression to test whether mtDNA mutations are correlated with latitude.

#### Statistical analyses of long-term and cross-sectional environmental data

##### Long-term monitoring and analyses of mound activity, number of reproductive offspring, and environmental variables between 2020-2023

Between 2020 and 2023, the activity of wood ant mounds, the emergence time and number of reproductive offspring (i.e. new queens and males), as well as the within-mound temperature were recorded from March to June in six populations Southern Finland. We observed three *F. aquilonia* populations (*Haltiala, Pusula, Soleböle*) and three *F. aquilonia* x *F. polyctena* hybrid populations (*Bunkkerikal, Langholmen, Svanvik*).

The within-mound temperature was measured using HOBO data loggers (HOBO 64K Pendant Temperature Data Logger, Bourne, USA). To analyse the within-mound temperature, we first calculated means for each month over the three observation years for hybrids and *F. aquilonia* separately. As the data was not normally distributed, we compared the means using Mann-Whitney-U tests.

The mound activity was recorded as a binomial (1 = active; 0 = non-active). We further recorded the date and counted and/or estimated the number of sexual offspring present on the mound (counted with 1-20 individuals; estimated with >20). For both mound activity and number of reproductive offspring, we calculated the relative percentages (i.e., relative proportions calculated over all years; count data for focal species/count data for both species). We used a proportion test for the mound activity to test whether the mound activity differed between hybrids and *F. aquilonia* for each month separately. We used a Pearson’s Chi-Squared test for count data to compare the number of estimated reproductives under the null hypothesis of no difference between the species for each month separately.

##### Analyses of cross-sectional environmental variables recorded during sampling

During sampling, we recorded the within-mound temperature and mound volume (detailed above). To assess whether hybrids and *F. aquilonia* differ in these variables in the sympatric area (i.e., latitude lower than 63.99), we compared the variables separately in a linear mixed effect model (LMER, lme4 package ^89^). The explanatory variables were the genomic assignment of taxa and the latitude of each sample, and the random factors were the population ID and the date of sampling. The model fit was checked using dHARMa ^90^. To calculate a r^2^ value for each model, we created an intercept-only model for each variable and included the same random effects. Specifically, we calculated the Nagelkerke Pseudo-r^2^ (squaredGLMM function, MuMIn package ^91^). Such a Pseudo-r^2^ value is a relative measure among similar models indicating how well the model explains the data and shows values for the fixed effects as well as fixed and random effects together. To check whether the species differed in the variables, we calculated pairwise contrasts (emmeans, adjusting for multiple testing; ^92^). We further tested whether the mound volume is associated with the latitude by correlating both in a linear regression – separately for hybrids and *F. aquilonia*. For this, we excluded one hybrid and one *F. aquilonia* outlier mounds with extremely large dimensions based on z-score outlier tests conducted for both taxa separately. We then split the hybrid and *F. aquilonia* data sets into a “southern” and “northern” area based on their respective latitudinal means and compared the means within taxa using a Welch’s t-test for hybrids and a Mann-Whitney-U-test for *F. aquilonia* (data not normally distributed). Using one-sided Mann-Whitney-U tests, we also tested whether hybrid and *F. aquilonia* mounds differ overall in the sympatric range and across entire Finland (for *F. aquilonia* mounds.

#### Critical Thermal Maximum (CT_max_) assay and statistical analyses

##### Conducting the CT_max_ assay

From each mound, four workers were used in an CT_max_ assay using a dry bath (Eppendorf block heater, Thermomixer R, Hamburg, Germany). We chose a dry bath because body size does not influence heat tolerance in a dry bath assay compared to using a hot plate ^49,93^. To avoid potential experimental acclimation, we tested the ants on the same day as they were sampled ^94^. Workers were transferred individually to an Eppendorf tube and sealed with cotton briefly before the start of the experiment to minimise the time each worker spent inside the tube. The tubes with workers were then placed in a block heater, which marked the start of the experiment. Workers were exposed to 40 °C for four minutes (Fig. 3 F inset). The temperature was then gradually increased to 42, 45, 47, and finally 49 °C. Each temperature step was held for approximately three minutes. Between temperature steps, the temperature of the block heater increased to the respective temperature in one or two minutes. These steps were selected based on pilot studies. In total, the assay lasted 20 minutes. As a control, we transferred one worker from each population individually to a single tube, sealed it with cotton, and kept it at room temperature to ensure that workers do not fall into a coma and die due to other circumstances.

Each minute, the state of the workers inside the tubes was recorded, with workers either being “active” or “knocked down”. This procedure was repeated until four workers from each mound were tested. A digital thermometer (“Prima”, Amarell, Kreuzwertheim, Germany) was placed inside a tube and sealed with cotton to measure the temperature inside the tube during the assay. This temperature was recorded at the beginning of each minute. The order in which the workers were placed in the dry bath was block-randomised to minimise potential heating differences of the block heater. To reduce any batch effects, at least one worker from each colony was tested in the same CT_max_ run.

##### Survival analysis and statistical associations with mtDNA and hybrid index

To check if ants from hybrid and *F. aquilonia* populations differ in the CT_max_ assay, we calculated proportion hazards of ants falling into heat coma using a Cox regression model (coxph function; survival package, ^95^). Such a Cox regression model allows using multiple fixed factors and a random factor. We used the species/hybrid identity, the parental mtDNA status (*F. polyctena* or *F. aquilonia* for hybrids; only *F. aquilonia* for *F. aquilonia*), the hybrid index, as well as weather data prior to sampling (calculations detailed below). We scaled all continuous data (the hybrid index and weather data) to make them comparable and used the latitude as random factor using the “frailty” function (survival package).

We calculated weather data prior to sampling for each mound by extracting temperature and precipitation data 21 days prior to sampling until the day of sampling from the Finnish Meteorological Institute data. We calculated the mean, minimum, maximum, and range (max - min) temperature and precipitation of 21 days including the day of sampling, 14 days + day of sampling, 7 days + day of sampling, 5 days + day of sampling, 3 days + day of sampling, one day before sampling, and the day of sampling. As some variables are correlated, we excluded highly correlated values using the function “findCorrelation”, which keeps the most uncorrelated representative variables (caret package; spearman-method, pairwise-complete.obs). From these variables, we kept the maximum and minimum temperature 21 days prior to sampling including the day of sampling, the minimum temperature 5 days prior to sampling, temperature range 7 and 3 days prior to sampling, maximum precipitation values 1 day prior to sampling, and the precipitation range 7 days prior to sampling.

Further, we conducted a survival analysis. For this, the first timepoint when a worker was recorded as knocked-down was selected and analysed (surv function, survival package, ^95^; plotted with survfit2 function, ggsurvfit package ^96^). We conducted a non-parametric Kaplan-Meyer survival analysis with the state of the worker as a response variable, and the time and the species/hybrid identity as explanatory variables. We then extracted the survival probabilities for minutes 8-12 from the survival analysis. These minutes encompass temperature values around 45 °C, which represent a biologically relevant temperature, when *Formica* ants fall into heat coma (identified in a pilot study, data not shown). To account for climatic changes but still compare hybrid and *F. aquilonia*, we conducted two approaches: In the first approach, we only selected samples from a sympatric area (i.e., latitude lower than 63.9), and in the second approach, samples from entire Finland.

As we found hybrids that were sired from either parent, *F. aquilonia* or *F. polyctena*, we tested whether the temperature at which ants fall into heat coma differs between the maternal lineages using a one-sided Mann-Whitney-U test. Additionally, we tested whether the hybrid index is correlated with the temperature at which ants fall into heat coma using a linear regression.

#### Persistence of hybrids and their expansion identified by 5-year climatic averages

##### Number and persistence of hybrids and geographic extent of hybrids

In the current sampling 2023, we re-visited 42 out of 75 populations from the previous sampling ^30^ that were already known to assess the number and persistence of hybrids as well as the geographic extent of hybridisation. Whenever possible, workers from the same mound were sampled, or workers from mounds within a distance of 500 metres, when the focal known mound was abandoned. Based on the taxa identity (i.e., species or hybrid) retrieved in both studies, we checked if the taxa persisted or changed. We also calculated the number of *F. aquilonia x F. polyctena* hybrid samples in both data sets and checked the latitude between data sets where hybrid populations were found. To check whether the frequency of hybrid samples has increased over time and with latitude conducted a multinomial logistic regression. For this, we lumped all cold-adapted (*F. aquilonia, F. lugubris*) and all warm-adapted (*F. pratensis, F. rufa*) species together and used the taxa identity (i.e., hybrids as well as cold-and warm-adapted species) as the response variables and the time (as factor) and latitude (as continuous variable) as explanatory variables (interaction model; multinom function, nnet package ^97^). We manually calculated a Z-score (i.e., Wald Z), two-tailed z-test, and a Nagelkerke Pseudo-r^2^ value for the model values. To check the model fit, we compared the model’s AICc with the AIC of an intercept-only model.

##### Calculation and statistical analyses of the 5-year climatic averages

To calculate climatic changes over time for the sampling locations, we downloaded temperature and precipitation data from the Finnish Meteorological Institute (FMI; Aalto et al., 2016) and extracted the data for each mound from digital rasters (FMI ClimGrid gridded daily climatology) at a 10-km spatial resolution for up to six years before the sampling year. We then calculated averages over 5 years before the respective sampling year. For example, if a mound from population was sampled in 2023, the climate for this mound was averaged over the years from 2018 to 2022. If it was sampled in 2005, it was averaged over the years from 1999 to 2004, etc. We calculated six climatic variables, i) winter mean temperature, ii) spring mean temperature, iii) winter mean precipitation, iv) spring mean temperature, v) the number of days above 0 °C in winter, and vi) the number of days below 0 °C in spring. Winter and spring were defined as the months of December to February and the months of April and May, respectively, and identically calculated as in Satokangas et al. (2023; ^30^) for comparability.

For the climatic change analysis, we used the current sampling as well as only samples from 2005 originating from the previous sampling. We then defined three ranges. A ‘hybrid’ range where hybrids were found in 2005, an ‘expanded’ range at higher latitudes where we found hybrids in 2023, and an ‘expanded’ range at higher latitudes where we did not find hybrids in 2005. Our rationale was as follows: If climatic changes have allowed a northward expansion of hybrids, we expect that the climate in the ‘expanded’ range has changed in a way that is favourable for hybrids. Specifically, we expect that the ‘expanded’ range in 2023, where hybrids have been found, should have a similar climate compared to that of the hybrid range in 2005.

We then conducted three comparisons. i) To test whether the climate has changed and allowed a northward expansion of hybrids, we compared the climate of hybrid populations in the ‘hybrid’ range found in 2005 (lower latitudes, range = 59.92 to 62.56, Fig. 3 B, light purple colour; N=5; ^30^) with the climate of hybrid populations in the ‘expanded’ range in 2023 at higher latitudes (latitudinal range = 62.57 to 63.97, Fig. 3 B, dark purple colour; N=4). ii) We also compared the climate of hybrid populations in the ‘hybrid’ range found in 2005 (lower latitudes, range = 59.92 to 62.56, Fig. 3 B, light purple colour) with the climate of hybrid populations in the ‘expanded’ range but using climatic data from 2005 where hybrids were absent (latitudinal range = 62.57 to 63.97, Fig. 3 B, yellow colour; N=4). iii) Lastly, we compared the climate of hybrid populations in the ‘expanded’ range in 2023 (higher latitudes, range = 62.57 to 63.97, Fig. 3 B, dark purple colour) with the climate of hybrid populations with the climate of hybrid populations in the ‘expanded’ range but using climatic data from 2005 (latitudinal range = 62.57 to 63.97, Fig. 3 B, yellow colour). Hybrids with different mtDNA parental species were combined in the comparison because they did not differ in the environmental variables.

Due to a low sample size in each range (N=4-5), we conducted a non-parametric bootstrap approach. For each comparison, we re-sampled the data 1,000 with replacement and calculated the difference in the means. To calculate a p-value, we extracted the 95% confidence intervals from the mean differences and counted the bootstrap values in which the mean difference was on the opposite side of zero from the observed mean, which indicates a significant difference between the data. We corrected the results for multiple testing by applying a Holm-Bonferroni correction.

We further tested whether the climate has overall changed in both hybrid and *F. aquilonia* populations over 18 years. For this, we compared the same six environmental variables described above within and between hybrid and *F. aquilonia* populations and between previous and current sampling. We compared the environmental variables using t-tests (if normally distributed) or Mann-Whitney-U tests (if not-normally distributed) and corrected the results for multiple testing.

