## Supplementary figures and tables for "Evidence for climate-mediated range expansion of hybrid wood ants"

**This PDF file includes:**

Figs. S1 to S7
Tables S1 to S8
Data S1 to S2

**Other Supplementary Materials for this manuscript include the following:**
Data S1 to S2
Photograph S1.

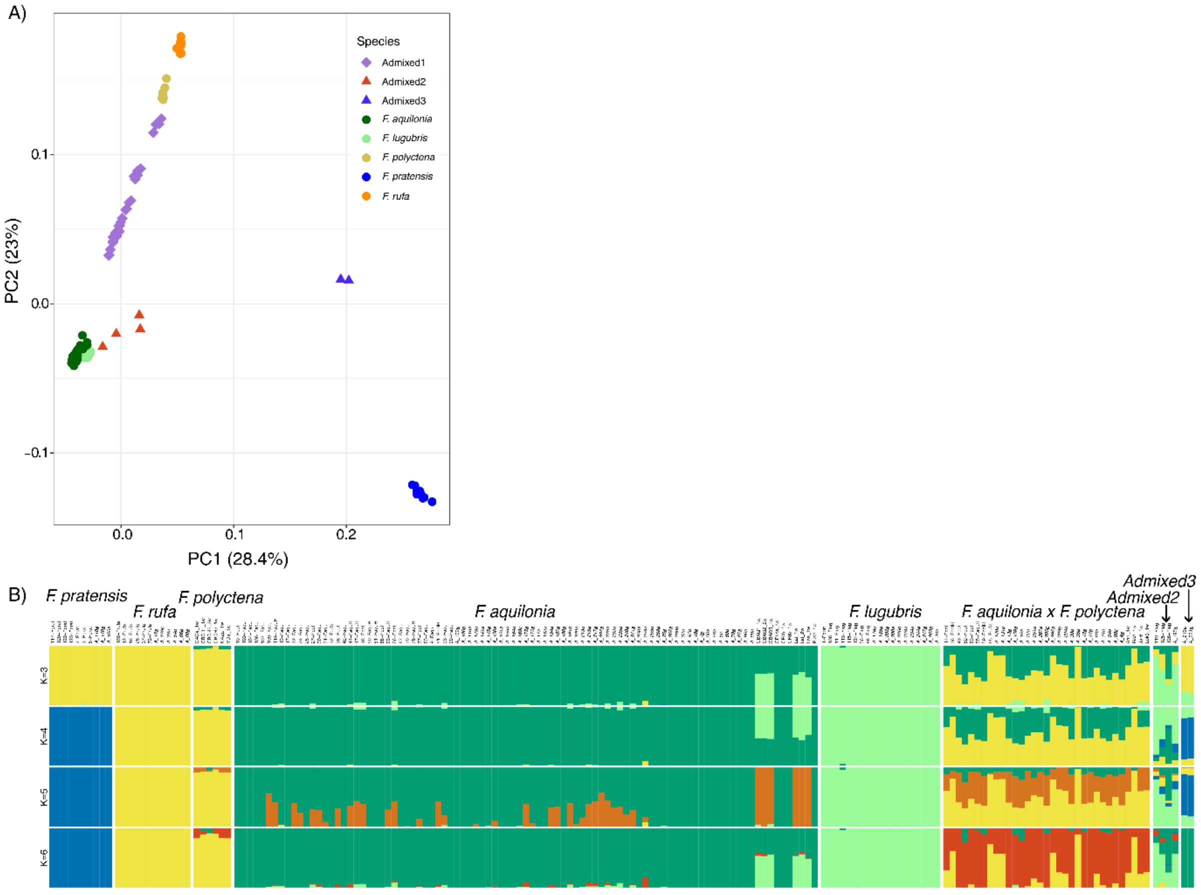


**Fig. S1. Downstream analyses of the whole-genome data for the five wood ant species and hybrids.** A) PCA with species assignment. In the PCA, *F. pratensis* samples clustered separately from other species and hybrid samples on the first axis (28.4%); only Admixed3 samples clustered nearby and were also separated from the other samples. The second axis (23%) separated mainly cold-adapted species *F. lugubris and F. aquilonia* from warm-adapted species *F. polyctena* and *F. rufa* and hybrid samples. Cold-adapted species *F. lugubris and F. aquilonia* clustered closely together and Admixed2 samples also clustered nearby to them, although more loosely aggregated. Warm-adapted species *F. polyctena* and *F. rufa* also clustered closely together but separately for each species, while hybrid (*F. aquilonia x F. polyctena*) samples clustered loosely between *F. aquilonia* and *F. lugubris* as well as *F. polyctena* and *F. rufa*. B) In the admixture plot, samples were identified as mainly 'pure' species and admixed taxa, and the same assignment of samples was retrieved as in the PCA.

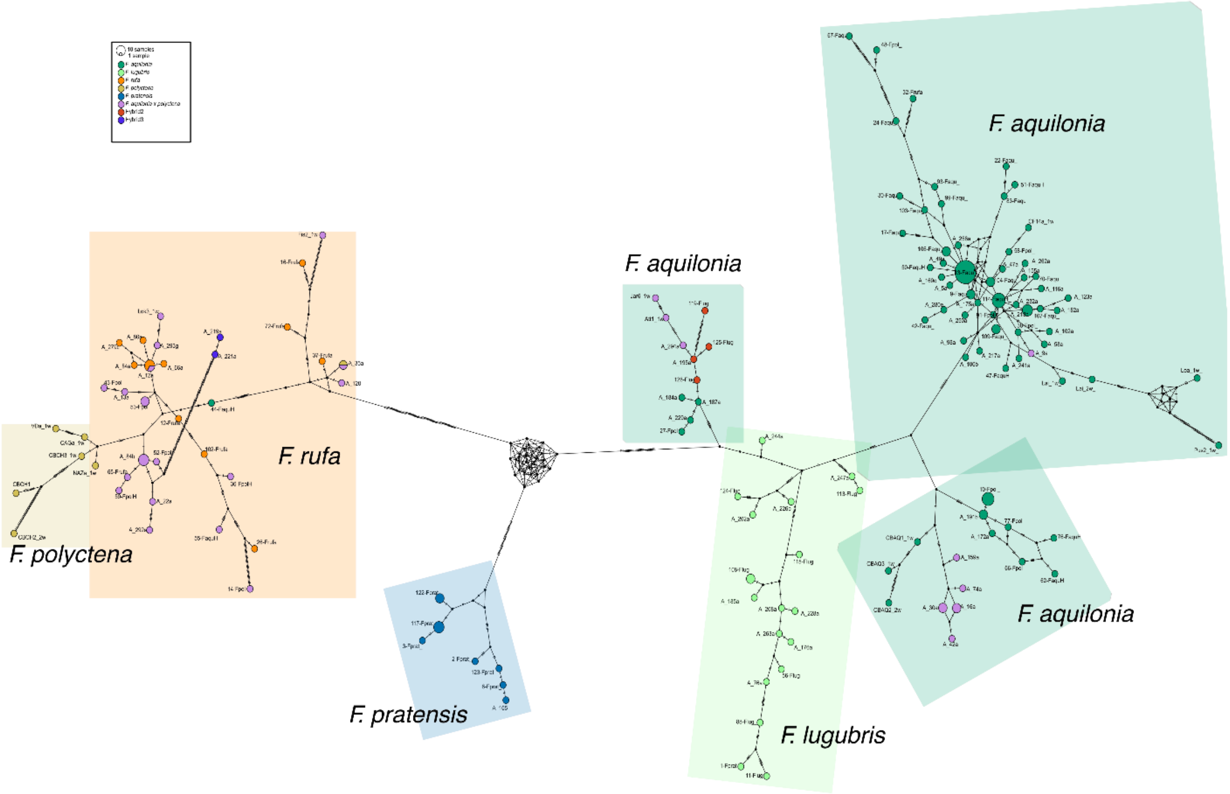


**Fig. S2. Mitochondrial haplotype network.** A minimum spanning network was created in PopArt) using all 2023 and reference samples ($N_{indsTotal}$=179, $N_{SNPs}$=466). Dot sizes represent the frequency of specific haplotypes- Dot colours represent the mtDNA origin, while coloured areas around the dots represent species identity based on whole-genome data $N_{SNPs}$=9979).

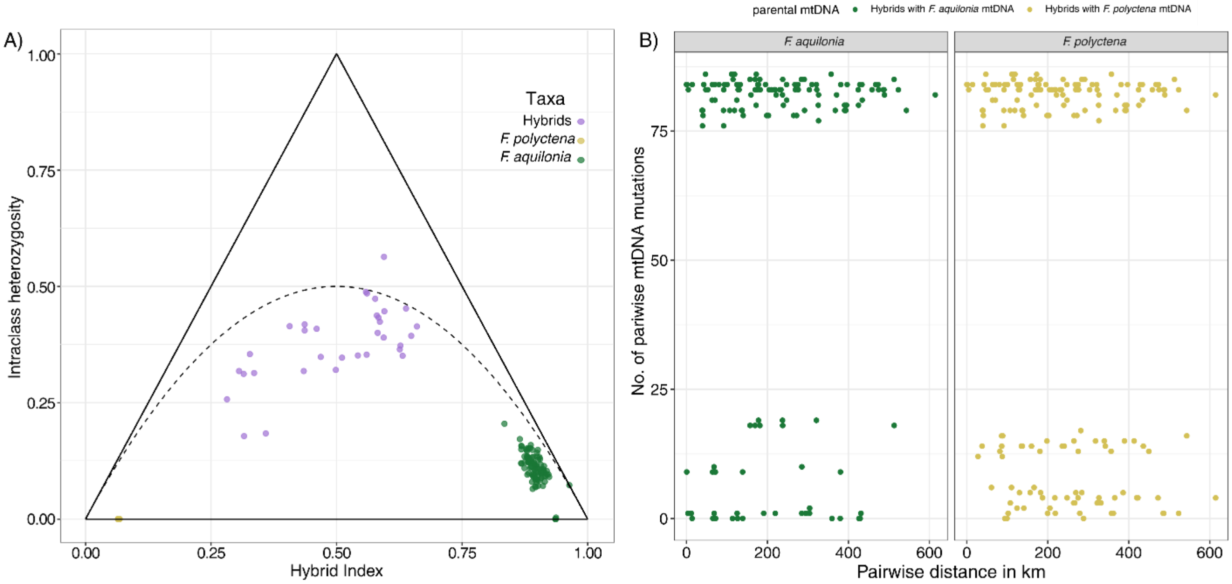


**Fig. S3. The A) hybrid index and B) number of mutations between hybrid sample pairs.**

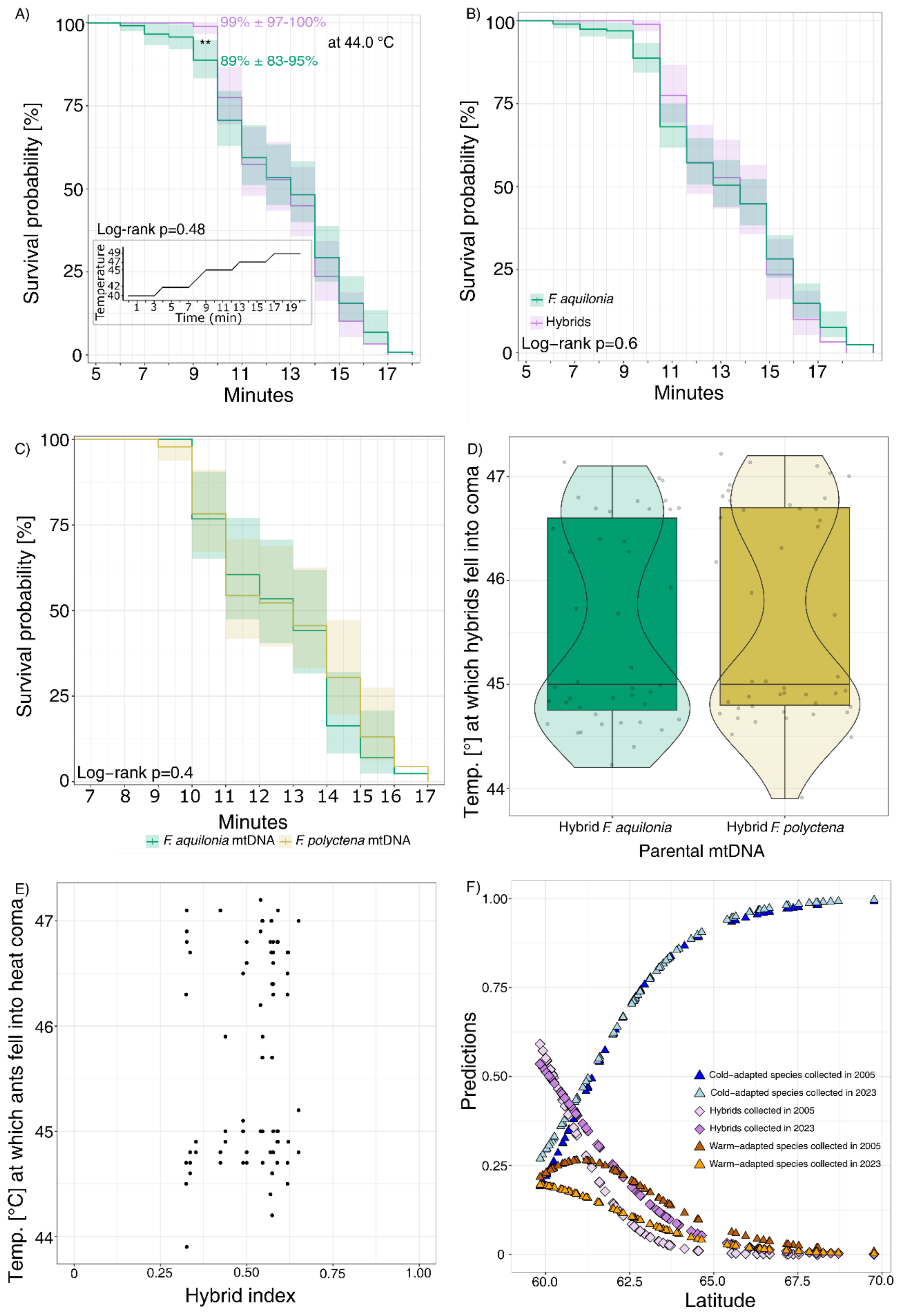
A)
99% ± 97-100%
**
at 44.0 °C
89% ± 83-95%
Survival probability [%]
Log−rank p=0.48
Temperature
Time (min)
Minutes
B)
Survival probability [%]
F. aquilonia
Hybrids
Log−rank p=0.6
Minutes
C)
Survival probability [%]
Log−rank p=0.4
Minutes
F. aquilonia mtDNA
F. polyctena mtDNA
D)
Temp. [°] at which hybrids fell into coma
Hybrid F. aquilonia
Hybrid F. polyctena
Parental mtDNA
E)
Temp. [°C] at which ants fell into heat coma
Hybrid index
F)
Predictions
Cold−adapted species collected in 2005
Cold−adapted species collected in 2023
Hybrids collected in 2005
Hybrids collected in 2023
Warm−adapted species collected in 2005
Warm−adapted species collected in 2023
Latitude

37

**Fig. S4. $CT_{max}$ assay results between *F. aquilonia* and hybrids and detailed analysis of hybris as well as the plot of the multinomial logistic regression.** A) Survival probabilities from the survival analysis for hybrids and F. aquilonia (workers tested, N hybrids = 89; N F. aquilonia = 116). The asterisks represent differences between the survival probabilities at minute 9 (43.9 °C; test of given proportions) with respective survival probabilities for hybrids and F. aquilonia. The inset in A) displays the $CT_{max}$ temperature increase over 20 minutes. B) No significant difference in the survival analysis between hybrids at *F. aquilonia* in entire Finland. C) No significant difference in the survival analysis between hybrids that originated from either *F. aquilonia* or *F. polyctena* maternal lineages (maternal mtDNA). D) Hybrids with *F. polyctena* mtDNA fall into heat coma at a higher temperature than hybrids with *F. aquilonia* mtDNA. E) No correlation between the hybrid index and the temperature when ants fall into heat coma. F) Predictions of the multinomial logistic regression to find warm- and cold-adapted species and hybrid populations at higher latitudes when comparing the 'previous' sampling conducted in 2005 with the 'current' sampling conducted in 2023.

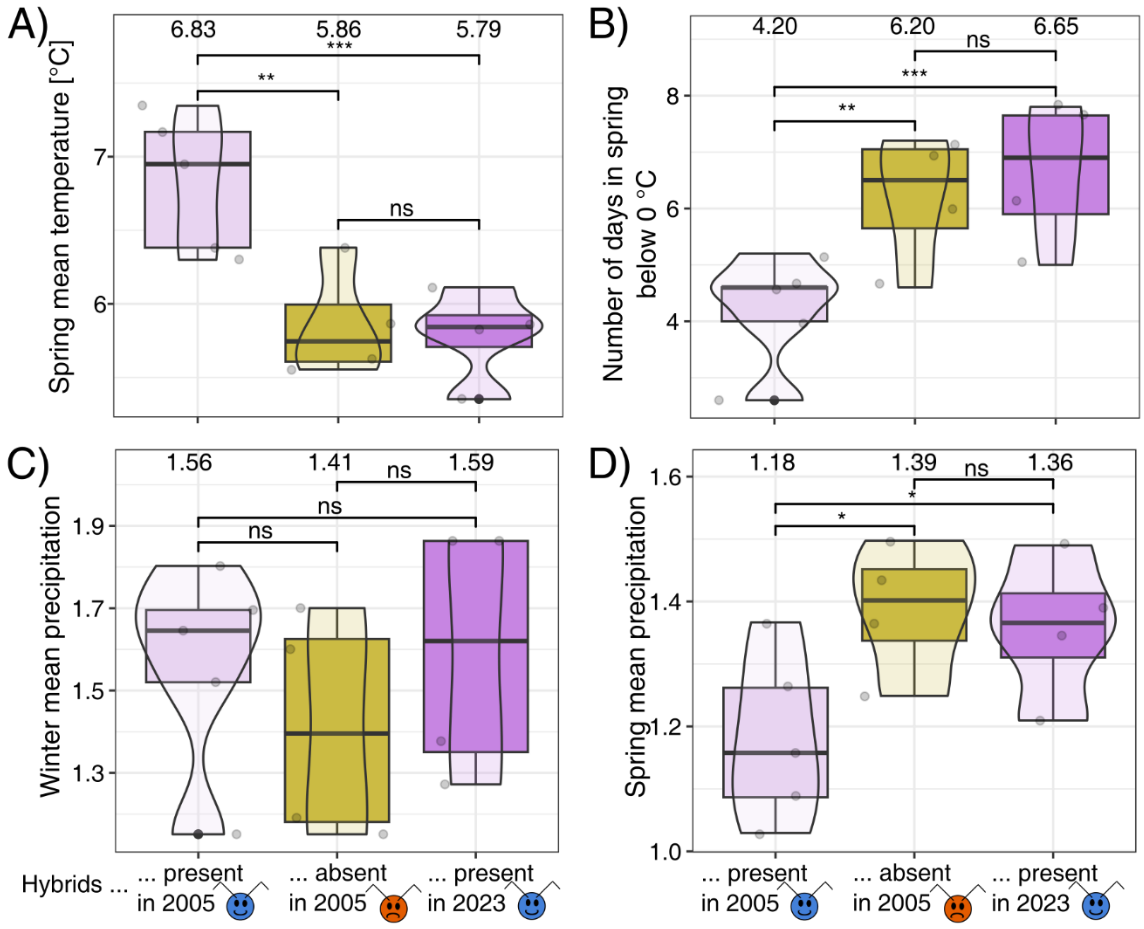


**Fig. S5. Climatic 5-year averages of four environmental variables are plotted for the hybrid populations only.** The variables were calculated backward from the sampling year 2005 (period 1999-2004) and 2023 (period 2018-2022). A) Spring mean temperature. B) Number of days in spring below 0 °C (frosty spring days). C) Winter mean precipitation. E) Spring mean precipitation. 'Winter' is defined as the diapause period between December to February and 'Spring' as the reproductive period between April to May. Values above the boxplots represent mean values. Asterisks represent bootstrapped p-values between hybrid areas at higher and lower latitudes and over time (Tab. S5) with * <0.05, ** <0.01, and ***<0.001. Please note that, we use the terms '2023' and '2005' but refer to the 5-year climatic average of the years 2018-2022 and 1999-2004, respectively. N suitable in 2005 = 5. N unsuitable in 2005 = 4. N suitable in 2023 = 4.

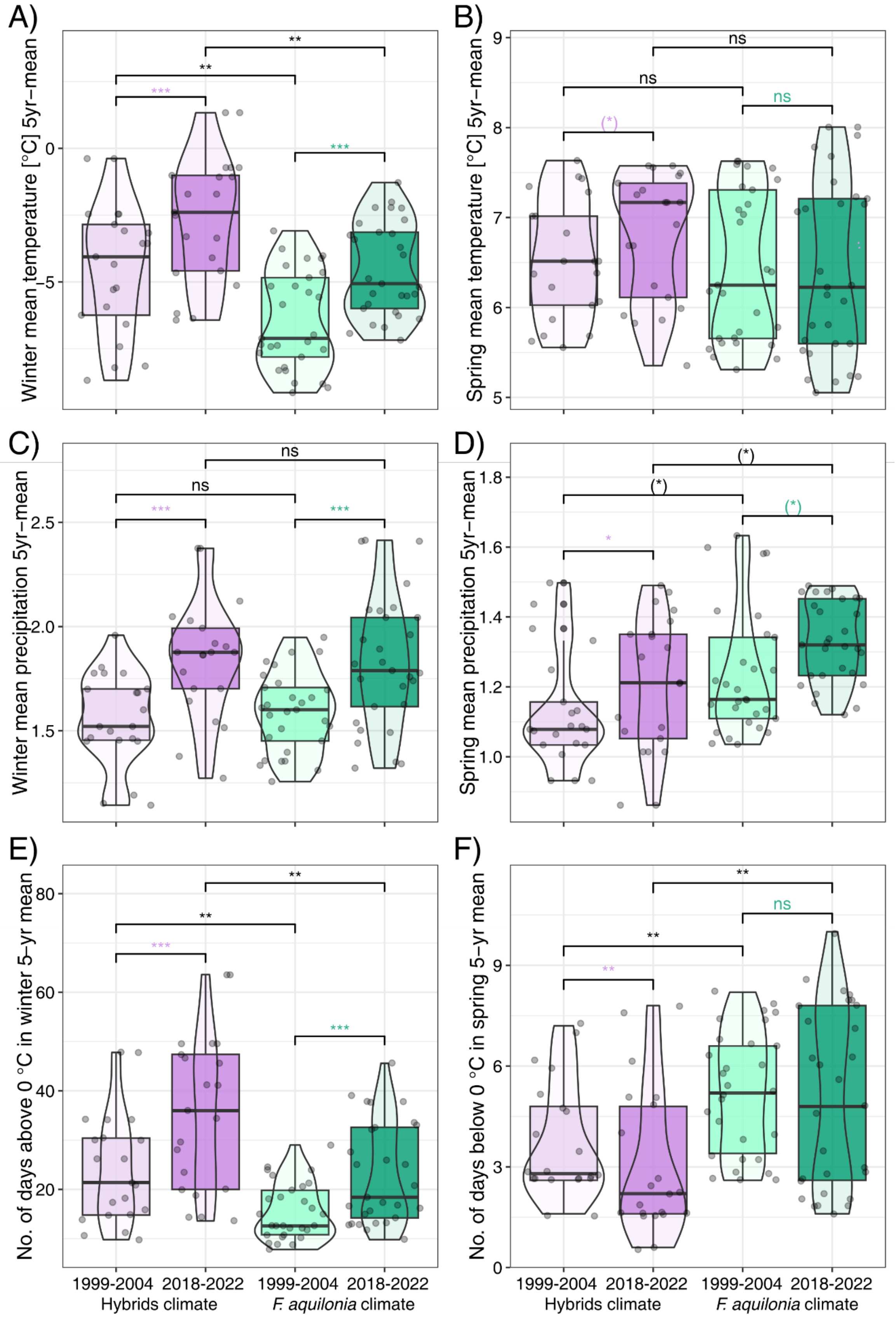

A)
B)
C)
D)
E)
F)
Winter mean temperature [°C] 5yr–mean
Spring mean temperature [°C] 5yr–mean
Winter mean precipitation 5yr–mean
Spring mean precipitation 5yr–mean
No. of days above 0 °C in winter 5–yr mean
No. of days below 0 °C in spring 5–yr mean
1999-2004 2018-2022 1999-2004 2018-2022
Hybrids climate
*F. aquilonia* climate


68

**Fig. S6. Environmental variables averaged over five years before the years 2005 (1999-2004) and 2023 (2018-2022) for the same locations.** A) Winter mean temperature, B) spring mean temperature, C) winter mean precipitation, D) spring mean precipitation, E) number of days above 0 °C in winter, F) number of days below 0 °C in spring. Purple and green colours represent hybrids and *F. aquilonia*, respectively. Comparisons within hybrids are coloured with purple asterisks and within *F. aquilonia* with green asterisks. Parentheses around asterisks indicate non-significance after correction for multiple testing. $N_{hybrids} = 21$. $N_{F.\ aquilonia} = 29$. If the climate has changed over 18 years, we expect to observe similar climatic changes in the sympatric area in both hybrid and *F. aquilonia* populations in key climatic variables. The winter of the years 2018-2023 was warmer, had more precipitation, and more days above 0 °C than the winter of the years 2000-2005 in both hybrid and *F. aquilonia* populations (coloured asterisks). Also, the spring of 2018-2023 had more precipitation and fewer frost days below 0 °C in hybrid but not *F. aquilonia* populations than the spring of the years 2000-2005 (coloured asterisks). Additionally, in hybrid populations, the winter was warmer and had more days above 0 °C, while the spring had fewer frost days below 0 °C compared to *F. aquilonia* populations in both 5-year average periods (all other variables did not differ between years and/or populations).

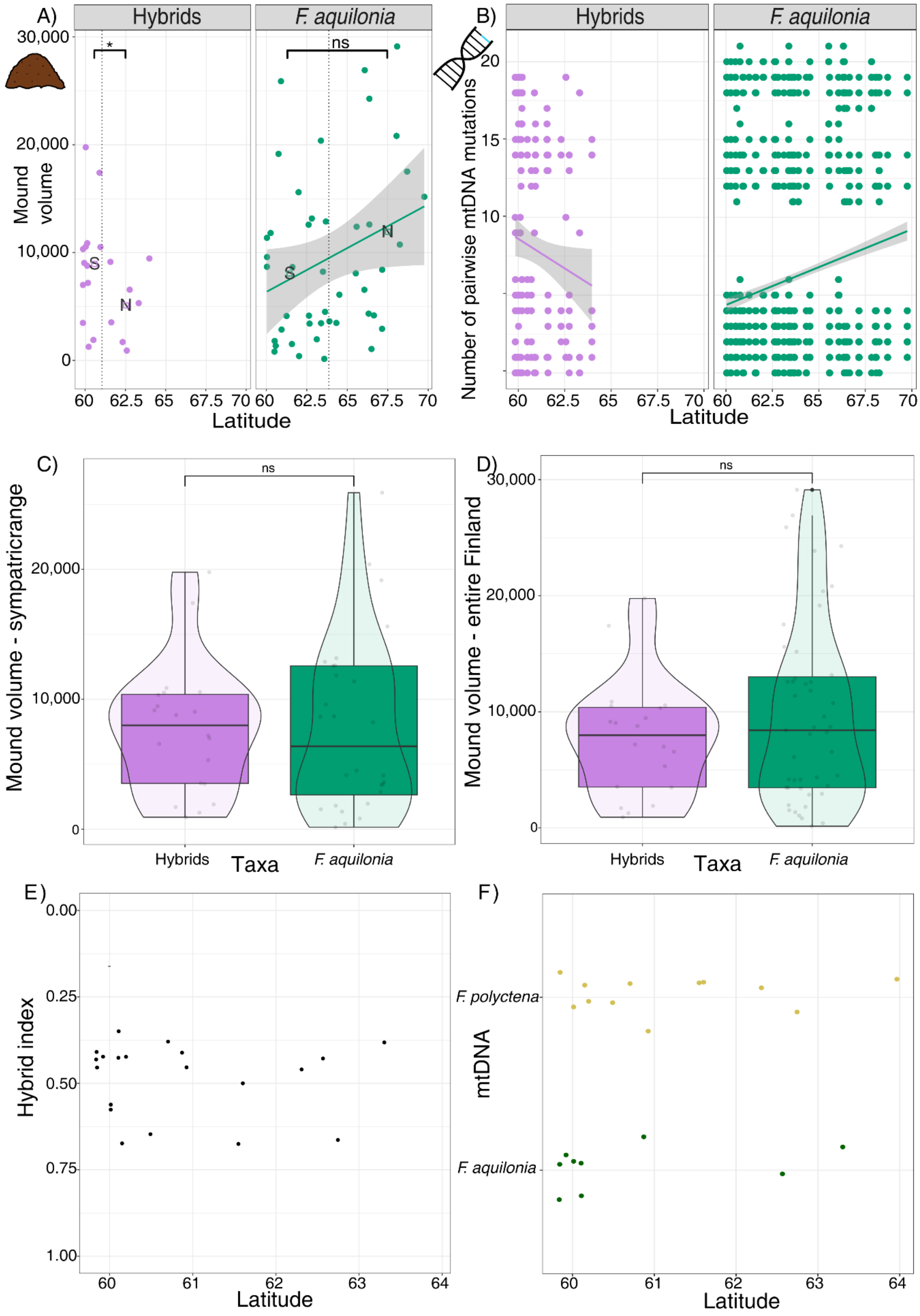
A)
Hybrids
F. aquilonia
30,000
20,000
10,000
0
Mound volume
*
ns
S
N
60
62.5
65
67.5
70
Latitude
B)
20
15
10
Number of pairwise mtDNA mutations
C)
ns
20,000
10,000
Mound volume - sympatricrange
Hybrids
Taxa
F. aquilonia
D)
Mound volume - entire Finland
E)
0.00
0.25
0.50
0.75
1.00
Hybrid index
61
62
63
64
F)
F. polyctena
F. aquilonia
mtDNA

87

**Fig. S7. Hybrid index and mtDNA origin plotted against latitude.** A) Calculated mound volume (based on mound length, width, and height) plotted against latitude for hybrids (N=21) and *F. aquilonia* (N=48). The dotted lines represent the latitudinal means for both hybrids and *F. aquilonia*. They split the data in a 'Northern' and 'Southern' area used to compare the means, which are represented as squares. Squares with 'S' and 'N' denote the means for the ''Southern' and 'Northern' area, respectively. C) The mound volume did not differ between hybrids and *F. aquilonia* in the sypatric range (means, hybrids: 8993.60; *F. aquilonia*: 10,948.11; one-sided Mann-Whitney-U-test, W = 466, p-value = 0.6274) or D) when tested across entire Finland (means, hybrids: 8993.60; *F. aquilonia*: 9892.90; one-sided Mann-Whitney-U-test, W = 312, p-value = 0.892). F) Number of pairwise mutations within hybrids and F. aquilonia separately plotted against the latitude for hybrids (N=572) and F. aquilonia (N=7310). In B), we combined data from the 'current' and 'previous' sampling to increase statistical power. We did not find a correlation between E) hybrid index and latitude or F) mtDNA origin (either *F. aquilonia* or *F. polyctena*) and latitude.

**Table S1. Mound GPS coordinates and characteristics recorded during fieldwork.**

| Internal ID | Population | Latitude | Longitude | Mound volume | Genomic ID | mtDNA ID |
|---|---|---|---|---|---|---|
| H294 | Aland | 60.013 | 20 | 10551 | Hybrids | *F. aquilonia* |
| H293 | Aland | 60.013 | 20 | 19773 | Hybrids | *F. polyctena* |
| H292 | Raisio | 60.491 | 22.148 | 1902 | Hybrids | *F. polyctena* |
| H285 | Ryjäojankulma | 60.871 | 22.271 | 17410 | Hybrids | *F. aquilonia* |
| H280 | Pirttilahdenkulma | 60.755 | 22.501 | 19157 | *F. aquilonia* | *F. aquilonia* |
| H267 | Virrat | 62.308 | 23.65 | 1707 | Hybrids | *F. polyctena* |
| H262 | Loukkukoskentie | 63.385 | 24.312 | 3457 | *F. aquilonia* | *F. aquilonia* |
| H258 | Veteli | 63.419 | 23.921 | 62083 | *F. aquilonia* | *F. aquilonia* |
| H255 | Kola | 63.584 | 23.855 | 150 | *F. aquilonia* | *F. aquilonia* |
| H253 | Sievie | 63.899 | 24.499 | 3631 | *F. aquilonia* | *F. aquilonia* |
| H251 | Karsikas | 63.966 | 25.435 | 9459 | Hybrids | *F. polyctena* |
| H250 | Kestilä | 64.328 | 26.298 | 3478 | *F. aquilonia* | *F. aquilonia* |
| H241 | Ruana | 66.06 | 26.368 | 6557 | *F. aquilonia* | *F. aquilonia* |
| H240 | Tervakari | 67.154 | 24.979 | 2937 | *F. aquilonia* | *F. aquilonia* |
| H235 | Kolari | 67.164 | 23.57 | 8417 | *F. aquilonia* | *F. aquilonia* |
| H232 | Muonio | 68.046 | 23.532 | 20822 | *F. aquilonia* | *F. aquilonia* |
| H229 | Sirkka | 67.837 | 24.83 | 23858 | *F. aquilonia* | *F. aquilonia* |
| H224 | Kevo | 69.762 | 26.991 | 15178 | *F. aquilonia* | *F. aquilonia* |
| H217 | Inarijärventie | 68.703 | 27.454 | 17522 | *F. aquilonia* | *F. aquilonia* |

| | | | | | | |
|---|---|---|---|---|---|---|
| H215 | Lahtola | 68.25 | 27.244 | 10748 | *F. aquilonia* | *F. aquilonia* |
| H213 | Vuotso | 68.085 | 27.181 | 29120 | *F. aquilonia* | *F. aquilonia* |
| H199 | Rovaniemi | 66.501 | 25.796 | 1066 | *F. aquilonia* | *F. aquilonia* |
| H191 | Kaarti | 66.65 | 29.111 | 4176 | *F. aquilonia* | *F. aquilonia* |
| H188 | Oulanka Research Station forest | 66.37 | 29.308 | 24273 | *F. aquilonia* | *F. aquilonia* |
| H187 | Oulanka Research Station | 66.37 | 29.312 | 12612 | *F. aquilonia* | *F. aquilonia* |
| H184 | Kuusamo | 66.324 | 29.262 | 4345 | *F. aquilonia* | *F. aquilonia* |
| H182 | Visala | 66.082 | 28.523 | 26918 | *F. aquilonia* | *F. aquilonia* |
| H175 | Taivalkoski | 65.579 | 28.236 | 12403 | *F. aquilonia* | *F. aquilonia* |
| H172 | Rantala | 65.531 | 27.761 | 8083 | *F. aquilonia* | *F. aquilonia* |
| H169 | Joumala | 64.517 | 25.122 | 6103 | *F. aquilonia* | *F. aquilonia* |
| H164 | Koivula | 63.502 | 26.086 | 8233 | *F. aquilonia* | *F. aquilonia* |
| H159 | Heinämäki | 63.303 | 26.81 | 5318 | Hybrids | *F. aquilonia* |
| H155 | Syrjälä | 63.376 | 26.894 | 20383 | *F. aquilonia* | *F. aquilonia* |
| H151 | Lapamäki | 63.681 | 27.544 | 12881 | *F. aquilonia* | *F. aquilonia* |
| H150 | Luokkimäki | 63.632 | 28.559 | 4511 | *F. aquilonia* | *F. aquilonia* |
| H145 | Pihlajavaara | 63.119 | 29.392 | 1963 | *F. aquilonia* | *F. aquilonia* |
| H141 | Eno | 62.814 | 30.133 | 13168 | *F. aquilonia* | *F. aquilonia* |
| H131 | Jakokoski | 62.745 | 30.036 | 6563 | Hybrids | *F. polyctena* |
| H123 | Härkinvaara | 62.674 | 29.581 | 3419 | *F. aquilonia* | *F. aquilonia* |
| H120 | Parikkala | 61.549 | 29.499 | 9144 | Hybrids | *F. polyctena* |
| H116 | Purnujärvi | 61.249 | 29.119 | 4126 | *F. aquilonia* | *F. aquilonia* |
| H106 | Mänlahti | 60.518 | 27.338 | 1811 | *F. aquilonia* | *F. aquilonia* |

| | | | | | | |
|---|---|---|---|---|---|---|
| H102 | Paavalinkylä | 60.505 | 26.232 | 821 | *F. aquilonia* | *F. aquilonia* |
| H98 | Pusula | 60.58 | 24.038 | 1361 | *F. aquilonia* | *F. aquilonia* |
| H96 | Ojajärvi | 60.907 | 24.01 | 25892 | *F. aquilonia* | *F. aquilonia* |
| H90 | Katoski | 60.925 | 24.456 | 10513 | Hybrids | *F. polyctena* |
| H87 | Miehola | 60.943 | 25.143 | 2866 | *F. aquilonia* | *F. aquilonia* |
| H84 | Hiivola | 60.703 | 24.688 | 34103 | Hybrids | *F. polyctena* |
| H74 | Perälänmäki | 62.566 | 22.791 | 920 | Hybrids | *F. aquilonia* |
| H70 | Mantylä | 61.582 | 21.925 | 1524 | *F. aquilonia* | *F. aquilonia* |
| H63 | Väha Myrräjärvia | 61.602 | 25.052 | 3537 | Hybrids | *F. polyctena* |
| H60 | Takajärvi | 61.616 | 25.311 | 8659 | *F. aquilonia* | *F. aquilonia* |
| H58 | Kiviperä | 62.614 | 24.999 | 12592 | *F. aquilonia* | *F. aquilonia* |
| H56 | Pukara | 62.636 | 26.201 | 4156 | *F. aquilonia* | *F. aquilonia* |
| H49 | Mikkeli | 62.027 | 26.906 | 405 | *F. aquilonia* | *F. aquilonia* |
| H47 | Kaakoporu | 61.994 | 26.078 | 15600 | *F. aquilonia* | *F. aquilonia* |
| H44 | Bunkkerikal | 59.843 | 23.233 | 7003 | Hybrids | *F. aquilonia* |
| H42 | Svanvik | 59.838 | 23.169 | 3481 | Hybrids | *F. aquilonia* |
| H35 | Fiskorvägen | 60.15 | 23.556 | 7193 | Hybrids | *F. polyctena* |
| H32 | Grabbskogvägen | 60.038 | 23.379 | 11370 | *F. aquilonia* | *F. aquilonia* |
| H30 | Grundsund | 59.921 | 23.039 | 9039 | Hybrids | *F. aquilonia* |
| H25 | Solebole | 60.039 | 23.047 | 8691 | *F. aquilonia* | *F. aquilonia* |
| H22 | Langholmen | 59.849 | 23.25 | 10329 | Hybrids | *F. polyctena* |
| H17 | Karis | 60.045 | 23.663 | 9590 | *F. aquilonia* | *F. aquilonia* |
| H16 | Trappberg, Pickala | 60.109 | 24.275 | 10870 | Hybrids | *F. aquilonia* |

| | | | | | | |
|---|---|---|---|---|---|---|
| H13 | Helsinki East | 60.197 | 25.108 | 1269 | Hybrids | *F. polyctena* |
| H9 | Marianne, Pickala | 60.105 | 24.266 | 8783 | Hybrids | *F. aquilonia* |
| H5 | Haltiala_A | 60.267 | 24.934 | 11820 | *F. aquilonia* | *F. aquilonia* |
| H4 | Haltiala_B | 60.261 | 24.935 | 12573 | *F. aquilonia* | *F. aquilonia* |

All samples were collected in Finland.

**Table S2. Number (and percentages) of assigned species and hybrid samples collected across Finland**

| Taxa | 'Previous' sampling 2005-2019 | 'Previous' and 'current' sampling - old and new populations | 'Current' sampling from 2023 – revisited populations | 'Current' sampling from 2023 – only new populations |
|---|---|---|---|---|
| *F. aquilonia* | 45 (52%) | 48 (52%) | 22 (50%) | 26 (54%) |
| *F. lugubris* | 8 (9%) | 11 (12%) | 4 (9%) | 7 (15%) |
| *F. polyctena* | 6 (7%) | 0 (0%) | 0 (0%) | 0 (0%) |
| *F. pratensis* | 7 (8%) | 3 (3%) | 2 (5%) | 1 (2%) |
| *F. rufa* | 6 (7%) | 6 (7%) | 4 (9%) | 2 (4%) |
| *F. aquilonia* x *F. polyctena* hybrids ('Admixed1' in (*28*) | 12 (14%) | 21 (23%) | 11 (25%) | 10 (21%) |
| *F. aquilonia* x *F. lugubris x F. rufa x F. pratensis* ('Admixed2' in (*28*) | 3 (3%) | 1 (1%) | 1 (2%) | 0 (0%) |
| *F. lugubris x F. polyctena/rufa* | 0 (0%) | 2[#] (2%) | 0 (0%) | 2 (4%) |
| **Total numbers** | **87** | **92** | **44** | **48** |

Note: # represents a newly detected hybrid combination. The 'previous' sampling (*28*) was collected mainly in 2005-2008 with additional samples collected until 2019. The 'current' sampling uses samples collected in 2023.

**Table S3. Mean within-mound temperature values measured from 2020 to 2023 for each month and for hybrid and *F. aquilonia* populations**

| Month of the year | Mean values hybrids | Mean values *F. aquilonia* | Mann-Whitney-U W | Mann-Whitney-U p-value | p-value corrected for multiple testing |
|---|---|---|---|---|---|
| 01 | 2.12 | 1.40 | 179903.0 | <0.001 | *** |
| 02 | 0.96 | 0.01 | 179534.5 | <0.001 | *** |
| 03 | 2.07 | 0.57 | 244460.5 | <0.001 | *** |
| 04 | 6.94 | 4.26 | 218912.5 | <0.001 | *** |
| 05 | 14.68 | 13.41 | 257571.5 | <0.001 | ** |
| 06 | 22.49 | 22.17 | 232133.5 | 0.218 | ns |
| 07 | 23.23 | 24.19 | 202922.5 | <0.001 | *** |
| 08 | 23.22 | 23.93 | 154003.5 | <0.001 | ** |
| 09 | 16.46 | 15.41 | 64365.0 | 0.001 | ** |
| 10 | 12.44 | 11.45 | 13015.5 | 0.004 | * |
| 11 | 10.23 | 8.52 | 377.0 | 0.269 | ns |
| 12 | 5.38 | 4.58 | 45878.0 | <0.001 | *** |

Significant comparisons are represented with asterisks, where * denotes p-value < 0.05, ** denotes p-value < 0.01, and *** denotes p-value < 0.001. ns = not significant.

**Table S4. Summary of the proportional hazard ratios.**

| Variable | Coefficient | Standard error of the coeffienct | Hazard ratio | 95% CI | p-value |
|---|---|---|---|---|---|
| **Species - *F. aquilonia*** | **1.04** | **0.47** | **2.82** | **1.11 - 7.13** | **0.028** |
| **mtDNA *F. polyctena*** | **-0.60** | **0.27** | **0.55** | **0.32 – 0.93** | **0.027** |
| **Hybrid index** | **-0.79** | **0.26** | **0.45** | **0.27 – 0.77** | **0.003** |
| Max. temperature 21 days prior to sampling | -0.22 | 0.17 | 0.80 | 0.58 – 1.11 | 0.182 |
| Min. temperature 21 days prior to sampling | 0.10 | 0.14 | 1.11 | 0.94 – 1.46 | 0.477 |
| Max. temperature 5 days prior to sampling | 0.29 | 0.17 | 1.33 | 0.96 – 1.85 | 0.088 |
| Temperature ranges 7 days prior to sampling | -0.10 | 0.11 | 0.91 | 0.73 – 1.23 | 0.382 |
| Temperature ranges 3 days prior to sampling | 0.21 | 0.11 | 1.23 | 0.99 – 1.53 | 0.066 |
| **Max. precipitation 1 day prior to sampling** | **0.34** | **0.10** | **1.40** | **1.14 – 1.71** | **0.001** |
| **Precipitation ranges 7 days prior to sampling** | **0.31** | **0.13** | **0.73** | **0.57 – 0.95** | **0.018** |

CI = Confidence intervals. Bold values represent statistically significant values.

**Table S5. Survival probability at measured temperatures and time points in the heat assay of hybrid and *F. aquilonia* populations in the sympatric area (i.e. overlapping geographic) and entire Finland**

| **Area** | **Temperature [°C] / time point when measured** | **Hybrid survival probability ± confidence intervals** | ***F. aquilonia* survival probability ± confidence intervals** | **Two-sample proportion test** |
|---|---|---|---|---|
| **Sympatric area** | 42.4 °C / 8 min | 1.00 ± 1.00-1.00 | 0.96 ± 0.92-1.00 | $X^2 = 2.30$ df = 1 p-value = 0.065 |
| | **44.0 °C / 9 min** | **0.99 ± 0.97-1.00** | **0.89 ± 0.83-0.95** | **$X^2 = 7.18$ df = 1 p-value = 0.004** |
| | 44.8 °C / 10 min | 0.78 ± 0.69-0.87 | 0.71 ± 0.63-0.80 | $X^2 = 0.95$ df = 1 p-value = 0.165 |
| | 44.7 °C / 11 min | 0.57 ± 0.47-0.68 | 0.60 ± 0.51-0.69 | $X^2 = 0.08$ df = 1 p-value = 0.613 |
| **Entire Finland** | 42.4 °C / 8 min | 1.00 ± 1.00-1.00 | 0.97 ± 0.95-0.99 | $X^2 = 1.35$ df = 1 p-value = 0.122 |
| | **43.9 °C / 9 min** | **0.99 ± 0.97-1.00** | **0.89 ± 0.84-0.93** | **$X^2 = 7.18$ df = 1 p-value = 0.004** |
| | 44.8 °C / 10 min | 0.78 ± 0.69-0.87 | 0.68 ± 0.62-0.75 | $X^2 = 2.06$ df = 1 p-value = 0.08 |
| | 44.7 °C / 11 min | 0.57 ± 0.47-0.68 | 0.57 ± 0.51-0.65 | $X^2 < 0.01$ df = 1 p-value = 0.500 |

Note: We selected four time points and their respective temperatures close to 45 °C, which is the temperature when most ants fell into heat coma. Note: Bold values represent statistically significant values based on confidence interval differences. df = degrees of freedom.

**Table S6. Observed species turnover of same mound or mound in near surroundings**

| Taxa turnover | Previous -> Now | Number of populations |
|---|---|---|
| *Formica* species to hybrids | *F. rufa* -> *F. aquilonia x F. polyctena* hybrid | 1 |
| | *F. lugubris* -> Admixed2<br>Admixed2 is a *F. aquilonia* x *F. lugubris x F. rufa x F. pratensis* hybrid indicating that the previous assignment to F. lugubris was due to a limited sample size | 1 |
| | *F. pratensis* -> *F. aquilonia x F. polyctena* hybrid | 1 |
| | *F. aquilonia* -> *F. aquilonia x F. polyctena* hybrid | 1 |
| Admixed2 to hybrid | Admixed2 -> *F. aquilonia x F. polyctena* hybrid (exact same mound) | 1 |
| hybrid to *Formica* species | *F. aquilonia x F. polyctena* hybrid<br>-> *F. rufa* | 1 |
| | *F. aquilonia x F. polyctena* hybrid<br>-> *F. aquilonia* | 2 |
| *Formica* species to other *Formica* species | *F. aquilonia* -> *F. rufa* (exact same mounds) | 2 |
| | *F. aquilonia* -> *F. lugubris* | 1 |
| | *F. lugubris* -> *F. aquilonia* (exact same mound) | 1 |

Note: In total, we detected 4 same mounds and 8 mounds in the surroundings with different species or hybrids

**Table S7. Comparing 5-year climatic averages (mean and bootstrapped tests) across ‘established and the ‘expanded’ ranges and between time**

| **Variable** | **Comparisons p-value / bootstrapped p-value** | **‘Established’ range with hybrids *present* in 2005 - lower latitudes (n=5)** | **‘Expanded range’ with hybrids *absent* in 2005 - higher latitudes (n=4)** |
|---|---|---|---|
| **Winter mean temperature** | **‘Expanded range’ with hybrids *absent* in 2005 - higher latitudes (n=4)** | 0.150 / **0.006**** | - |
| | **‘Expanded range’ with hybrids *present* in 2023 - higher latitudes (n=4)** | 0.730 / 0.442 | 0.230 / **0.006**** |
| **Spring mean temperature** | **‘Expanded range’ with hybrids *absent* in 2005 - higher latitudes (n=4)** | 0.098 / **0.004**** | - |
| | **‘Expanded range’ with hybrids *present* in 2023 - higher latitudes (n=4)** | **0.048 / <0.001***** | 0.886 / 0.840 |
| **Winter mean precipitation** | **‘Expanded range’ with hybrids *absent* in 2005 - higher latitudes (n=4)** | 1.00 / 0.792 | - |
| | **‘Expanded range’ with hybrids *present* in 2023 - higher latitudes (n=4)** | 1.00 / 0.916 | 1.00 / 0.792 |
| **Spring mean precipitation** | **‘Expanded range’ with hybrids *absent* in 2005 - higher latitudes (n=4)** | 0.260 / **0.012*** | - |

| | | | |
|---|---|---|---|
| | **‘Expanded range’ with hybrids *present* in 2023 - higher latitudes (n=4)** | 0.260 / **0.024*** | 0.690 / 0.666 |
| **Winter number of days above 0 °C** | **‘Expanded range’ with hybrids *absent* in 2005 - higher latitudes (n=4)** | 0.150 / **0.006**** | - |
| | **‘Expanded range’ with hybrids *present* in 2023 - higher latitudes (n=4)** | 0.340 / 0.130 | 0.340 / **0.032*** |
| **Spring number of days below 0 °C** | **‘Expanded range’ with hybrids *absent* in 2005 - higher latitudes (n=4)** | 0.120 / **0.004**** | - |
| | **‘Expanded range’ with hybrids *present* in 2023 - higher latitudes (n=4)** | 0.110 / **<0.001***** | 0.340 / 0.572 |

Note: Parentheses around a p-value indicate its non-significance after correction for multiple testing. Note, we use the terms ‘2023’ and ‘2005’ for simplicity but refer to the 5-year climatic average of the years 2018-2022 and 1999-2004, respectively. Significant comparisons are represented with asterisks, where * denotes p-value < 0.05, ** denotes p-value < 0.01, and *** denotes p-value < 0.001.

**Table S8. Comparing 5-year climatic averages between hybrid and *F. aquilonia* populations**

| **Variable** | **Year (5-year average)** | **Variable mean hybrids** | **Variable mean *F. aquilonia*** | **Test, test statistic, (df), p-value** |
|---|---|---|---|---|
| **Winter mean temperature** | 2005 (1999-2004) | -4.50 | -6.45 | **t-test, t = -3.09, df = 35.52, p-value = 0.004** |
| | 2023 (2018-2022) | -2.63 | -4.53 | **t-test, t = -3.14, df = 35.38, p-value = 0.004** |
| **Test, test statistic, df, p-value** | | **paired t-test, t = -49.44, df = 20, p-value<0.001** | **paired t-test, t = -64.39, df = 28, p-value<0.001** | |
| **Spring mean temperature** | 2005 (1999-2004) | 6.53 | 6.45 | Wilcox-test, W = 276, p-value = 0.582 |
| | 2023 (2018-2022) | 6.80 | 6.42 | Wilcox-test, W = 225, p-value = 0.120 |
| **Test, test statistic, df, p-value** | | paired Wilcox-test, V = 55, (p-value = 0.037) | paired Wilcox-test, V = 253, p-value = 0.449 | |
| **Winter mean precipitation** | 2005 (1999-2004) | 1.56 | 1.59 | t-test, t = 0.48, df = 39.10, p-value = 0.638 |
| | 2023 (2018-2022) | 1.84 | 1.83 | t-test, t = -0.17, df = 45.85, p-value = 0.866 |
| **Test, test statistic, df, p-value** | | **paired t-test, t = -6.43, df = 20, p-value<0.001** | **paired t-test, t = -5.35, df = 28, p-value<0.001** | |

| | | | | |
|---|---|---|---|---|
| **Spring mean precipitation** | 2005 (1999-2004) | 1.13 | 1.24 | **Wilcox-test,<br>W = 434,<br>p-value = 0.011** |
| | 2023 (2018-2022) | 1.21 | 1.33 | Wilcox-test,<br>W = 411,<br>(p-value = 0.037) |
| **Test, test statistic, df, p-value** | | paired Wilcox-test,<br>V = 54,<br>(p-value = 0.034) | paired Wilcox-test,<br>V = 117,<br>(p-value = 0.031) | |
| **Winter number of days above 0 °C** | 2005 (1999-2004) | 24.27 | 15.50 | **Wilcox-test,<br>W = 156,<br>p-value = 0.004** |
| | 2023 (2018-2022) | 35.74 | 23.12 | **Wilcox-test,<br>W = 149.5,<br>p-value = 0.002** |
| **Test, test statistic, df, p-value** | | **paired Wilcox-test,<br>V = 0, p-value<0.001** | **paired Wilcox-test,<br>V = 0,<br>p-value<0.001** | |
| **Spring number of days below 0 °C** | 2005 (1999-2004) | 3.73 | 5.21 | **Wilcox-test,<br>W = 446.5,<br>p-value = 0.005** |
| | 2023 (2018-2022) | 3.05 | 5.00 | **Wilcox-test,<br>W = 456.5,<br>p-value = 0.003** |
| **Test, test statistic, df, p-value** | | **paired Wilcox-test,<br>V = 205,<br>p-value = 0.002** | paired Wilcox-test,<br>V = 261.5,<br>p-value = 0.347 | |

Mean values and statistical results are denoted for the six environmental variables. Note: Parentheses around a p-value indicates its non-significance after correction for multiple testing. Significant values are in bold font. df = degrees of freedom.

**Data S1. (separate file) Data1.xlsx.** Excel file with all relevant data to re-analyse the main results.

**Data S2. (separate file) Wood_ants_Finland_MS.R.** R-script to re-analyse the main results.

**Photograph S1. (separate file) Photograph of a mound and its surface temperature in plain sunlight during summer related to the section "*Hybrids perform better under acute heat stress*".**
